# Cardiac Hemorrhage Precedes Hypertension-induced Fibrosis in Plasminogen Activator Inhibitor-1 Deficient Mice

**DOI:** 10.1101/2025.11.19.689269

**Authors:** Alex C. Pettey, Sohei Ito, Michael K. Franklin, Deborah A. Howatt, Jessica J. Moorleghen, Bryana M. Levitan, David B. Graf, Valerie Z. Guzmán, Nancy Zhang, Daniel A. Lawrence, Thomas H. Sisson, Hisashi Sawada, Jeffrey E. Saffitz, Hong S. Lu, Alan Daugherty

## Abstract

**Aims:** Plasminogen activator inhibitor-1 (PAI-1) regulates plasmin-mediated proteolysis, thereby influencing vascular stability and tissue remodeling. Angiotensin II (AngII) induces an increase in PAI-1 during the development of ascending thoracic aortic aneurysm (ATAA). The initial purpose of this study was to determine whether deletion of PAI-1 influenced development of ATAA. Subsequently, this study aimed to define the early pathological events preceding cardiac fibrosis in PAI-1 deficiency and the structural domain responsible for its protective effect.

**Methods and results:** AngII was infused for 4 weeks in whole-body PAI-1 deficient (PAI-1−/−) mice and their wild-type littermates (PAI-1+/+) to examine the role of PAI-1 in ATAA. PAI-1 deficiency did not alter AngII-induced aortopathy but revealed a striking cardiac phenotype characterized by replacement fibrosis predominantly within the epicardium and posterior septum. Ferric iron, indicative of prior hemorrhage, was coincident with fibrosis. Similar phenotypes were observed in PAI-1−/− mice infused with norepinephrine for 4 weeks. To define the pathological events preceding cardiac fibrosis, either AngII or norepinephrine was infused for 1 week in PAI-1+/+ or −/− mice. Both infusions induced extensive epicardial hemorrhage and posterior septal fibrosis in PAI-1−/− mice. To explore the initiation of cardiac pathology, AngII was infused for approximately 1 day. PAI-1−/− mice developed diffuse hemorrhage and cardiomyocyte injury localized to the posterior septum, pathologic changes that preceded overt fibrosis. Finally, to determine the protective domain of PAI-1, saline or AngII was administered to mice harboring loss-of-function point mutations in the protease inhibitory (PAI-1^Ala/Ala^) or somatomedin B-binding domains (PAI-1^AK/AK^). Compared to saline infusion, 1 week of AngII induced hemorrhage and heterogeneous fibrosis in PAI-1^Ala/Ala^, but not PAI-1^AK/AK^ mice.

**Conclusions:** These findings support that, under hemodynamic stress, PAI-1 deficiency promotes early cardiac hemorrhage and cardiomyocyte injury that lead to fibrosis. Mutational studies implicate dysregulated plasmin generation as an initiator of cardiac injury and fibrosis.

**TRANSLATIONAL PERSPECTIVE:** Cardiac fibrosis has been reported in a human population with PAI-1 deficiency and currently lacks targeted therapy. Our findings demonstrate that in animal models, PAI-1 deficiency confers susceptibility to cardiac injury in response to hemodynamic stress, which may accelerate fibrotic remodeling. Mutational disruption of the protease-inhibitory domain of PAI-1 induced similar pathology, supporting a protective role for this function. These observations suggest that interventions aimed at controlling hypertension, promoting endothelial integrity, or regulating plasmin activation could reduce fibrotic remodeling in this population.

## INTRODUCTION

Cardiovascular diseases (CVDs) are a leading cause of death globally and represent a major healthcare burden ^1^. Despite therapeutic advances, many forms of CVD lack effective treatments, particularly aortic aneurysms and cardiomyopathies ^2, 3^. This need has motivated continued investigation into the molecular mechanisms that contribute to cardiovascular pathologies. Plasminogen activator inhibitor-1 (PAI-1), a serine protease inhibitor, has been implicated in several CVDs and has received considerable attention as a potential therapeutic target ^4, 5^. In experimental models, genetic deficiency of PAI-1 in mice reduces fibrosis in most organs ^6^. In contrast, PAI-1 deficient mice develop spontaneous cardiac fibrosis with aging, and show exacerbated fibrosis in response to angiotensin II (AngII) infusion, either alone or in combination with aldosterone ^7–13^. PAI-1 has also been reported to limit the development of abdominal aortic aneurysms (AAAs) ^14–17^. Overall, observations in animal models suggest that PAI-1 exerts context-dependent protective or adverse effects in CVD.

PAI-1 is the primary inhibitor of tissue-type (tPA) and urokinase-type (uPA) plasminogen activators and thereby limits plasminogen activation, a function mediated by the reactive center loop (RCL) domain. PAI-1 also promotes lung fibrosis through its somatomedin B (SMB)-binding domain, independently of protease inhibition ^18, 19^. Through regulation of plasmin-mediated proteolysis, PAI-1 contributes to fibrin stability and influences extracellular matrix (ECM) turnover by restricting direct degradation of ECM or modulating activation of matrix metalloproteinases (MMPs) ^20^. These functions place PAI-1 at the interface of vascular integrity and tissue remodeling. Previous studies reported that pharmacological inhibition or genetic deficiency of PAI-1 attenuates AngII-induced aortic remodeling in mice ^8, 13^. Consistent with this finding, our recent study demonstrated that PAI-1 protein was increased markedly in the ascending aortas of AngII-infused mice and was abundant in human ascending thoracic aortic aneurysm (ATAA), implicating a potentially pathological role of PAI-1 in ATAA. While PAI-1 has been shown to protect against AAA in animal models ^14–17^, its role in ATAA, a disease process distinct from AAA, has not yet been reported.

To determine the role of PAI-1 in ATAA, whole-body PAI-1 deficient mice (PAI-1−/−) and their wild-type littermates (PAI-1+/+) were infused with AngII for 4 weeks. PAI-1 deficiency did not alter AngII-induced ATAA. Consistent with previous studies ^8, 11–13^, pronounced cardiac fibrosis was observed in AngII-infused PAI-1−/− mice. This phenotype was evident in both male and female mice. Prior reports have suggested that PAI-1 may protect against microvascular injury in the heart ^9, 12^, but the link between vascular integrity and cardiac fibrosis has not been defined. Moreover, the sequence of pathological events by which PAI-1 deficiency promotes cardiac fibrosis and the structural domain responsible for its protective effect remain uncertain. Therefore, the temporal and spatial characteristics of cardiac pathologies exacerbated by PAI-1 deficiency were determined. Further, norepinephrine (NE) was infused at a pressor rate to dissect the roles of hypertension versus other effects of AngII on development of cardiac pathologies in PAI-1−/− mice. Finally, mice harboring loss-of-function point mutations in the RCL and SMB-binding domain were generated and infused with AngII to delineate the structure-function roles of PAI-1.

## METHODS

### Mice

PAI-1+/− (B6.129S2-*Serpine1*^*tm1Mlg*^/J, Strain #002507) mice were purchased from The Jackson Laboratory. Littermate PAI-1+/+ and PAI-1−/− study mice were generated from male and female PAI-1+/− breeders. PAI-1^AK/AK^ and PAI-1^Ala/Ala^ study mice were developed as described previously^21^ and generated from breeders homozygous for their respective mutations. Mice were maintained on a 14-hour light/10-hour dark cycle at a controlled temperature of 20-23 °C and had free access to water and a standard laboratory diet (#2918, Inotiv). Inclusion and exclusion criteria for study mice were described in the Supplemental Materials following the ARRIVE Guidelines Checklist ^22^.

### Subcutaneous infusion using an osmotic mini pump

Mice (8 to 14-weeks of age) were infused in separate studies with saline, AngII, or NE through subcutaneously implanted osmotic mini pumps. AngII (1,000 ng/kg/min, #H-1705, Bachem) was dissolved in saline. NE (5.6 mg/kg/day, #A9512, MilliporeSigma) was dissolved in L-ascorbic acid (0.2% w/v in saline, #A65-21, Fisher Scientific) and protected from light. For 1 day or 1 week of infusion, osmotic mini pumps (Alzet model 1007D or 2001) were primed overnight at 37 °C in saline to initiate reagent release before pump implantation. For 4 weeks of infusion, osmotic mini pumps (Alzet model 1004) were primed overnight at 37 °C in saline to initiate infusion approximately 1 day after pump implantation, and surgical staples used to close incision sites were removed 7 days after surgery. Postoperative pain was alleviated by application of topical lidocaine cream (4% wt/wt, #59-930, HealthWise).

### Ultrasound measurements

Ultrasound images were captured using a Vevo 3100 system equipped with an MX550D (25–55 MHz) linear transducer (VisualSonics, Fujifilm). Mice were anesthetized using inhaled isoflurane (2-3% vol/vol) and maintained at a heart rate of >400 beats per minute during image capture to reduce anesthesia exposure and maintain consistent heart rates between animals (SomnoFlo, Kent Scientific). M-mode was obtained from the short axis bisecting the ventricle at the level of the papillary muscles. Three consecutive m-mode clips were obtained and 3 consecutive cardiac cycles from each m-mode clip were traced manually and averaged using Vevo LAB (v5.8.2, VisualSonics). The order in which mice were subject to ultrasound was not pre-selected. Image capture and data analysis were performed with animal genotype, infusion, and sex blinded.

Aortic imaging was performed using our standardized protocols ^23^. Briefly, images of the ascending aorta were captured using the right parasternal view and standardized according to two anatomical landmarks: the innominate artery branch point and the aortic valve. Aortic lumen was defined as the edge-to-edge distance of the ascending aortic wall between the sinotubular junction and the innominate artery. The largest luminal diameters of ascending aortas were measured in end-diastole over three cardiac cycles and averaged.

The heart was visualized from the modified parasternal long-axis and short-axis views. The left ventricular dimensions and calculated left ventricular ejection fraction were measured from the short-axis M-mode view ^24^. Left ventricular systolic strain was derived from Vevo LAB utilizing speckle tracking of the modified parasternal long-axis and short-axis images. Each mouse consisted of three technical replicates that were averaged.

### Quantification of aortic diameters using *in situ* images

Mice surviving 4 weeks of AngII infusion were euthanized by a cocktail of ketamine and xylazine (90 mg/kg #11695–6840-1, 10 mg/kg #11695–4024-1, respectively; Covetrus). The right atrial appendage was excised, and saline (~10 ml) was perfused via the left ventricle. Periaortic tissues were dissected carefully out, and a black plastic sheet was inserted underneath the aorta to improve image contrast ^25, 26^. A millimeter ruler was placed next to the aorta for calibration of measurements. *In situ* aortic images were captured with a Nikon SMZ25 stereoscope (Nikon). Aortic images were analyzed using NIS-Elements AR software (Version 5.11, Nikon). To measure aortic diameters, a measurement line was drawn perpendicularly to the aortic axis at the most dilated area of the ascending aorta. Measurements were verified by an individual who was blinded to the study groups.

### Histology and immunostaining

Samples were randomly selected for histology when no gross pathology was observed, or to represent the full range of observed gross pathology when present. For histological analyses, tissues were harvested from surviving mice, immersed in neutral-buffered formalin (10% w/v), incubated in ethanol (70% v/v), and subsequently embedded in paraffin. Paraffin-embedded sections (5 µm) were deparaffinized using limonene (#183164, MilliporeSigma). Masson’s trichrome (#ab150686, abcam) staining was performed according to the manufacturer’s protocol. For Verhoeff elastin staining, sections were stained with Verhoeff iron hematoxylin (hematoxylin 2.8% w/v, #3803815, Surgipath; ferric chloride 2.2% w/v, #3120-32, Ricca; iodine 0.4% w/v, #041955.22, Thermo Scientific; and potassium iodide 0.9% w/v, #0512, VWR; in 56% v/v ethanol) for 20 minutes and destained in 2% ferric chloride for 1-2 minutes. For Prussian blue staining, sections were stained with 1% w/v potassium ferrocyanide (#424135000, Thermo Scientific) in hydrochloric acid (0.37% v/v, #258148, MilliporeSigma) for 30 minutes and destained in neutral red (0.33% w/v, #N2889, MilliporeSigma) for 5 minutes. For picrosirius red, fast green, and Prussian blue co-staining, sections were stained with 1% w/v potassium ferrocyanide (#424135000, Thermo Scientific) in hydrochloric acid (0.37% v/v, #258148, MilliporeSigma) for 30 minutes followed by picrosirius red/fast green solution (0.1% w/v direct red, #365548, 0.1% w/v fast green, #F7252, in 1.3% w/v picric acid solution, #P6744, MilliporeSigma) for 60 minutes.

Immunostaining was performed using blocking reagents, primary, and secondary antibodies listed in the Supplementary Materials Major Resources Tables. Deparaffinized sections were incubated with H_2_O_2_ (1% v/v, #H325, Fisher Scientific; in methanol) for 2 minutes at 40 °C. Antigen retrieval was performed using EZ-AR antigen retrieval solution 2 (#HX032YCX-GP, BioGenex) at 98 °C for 20 minutes and cooled at 20-25 °C for 10 minutes. Nonspecific binding sites were blocked for 20 minutes at 20-25 °C. Primary antibodies were incubated overnight at 4 °C. Secondary antibodies were incubated at 20-25 °C for 30 minutes. Horseradish peroxidase-conjugated antibodies were visualized using NovaRED chromogen (#SK-4805, Vector). Coverslips were applied for histology and IHC stains using Permount (#SP-15, Fisher Scientific). For co-IF stains, autofluorescence was quenched using TrueVIEW (#SP-8400-15, Vector), and coverslips were applied using VECTASHIELD mounting medium with DAPI (#H-1200-10, Vector). Images of histological staining and immunostaining were captured using an Axioscan Z1 or 7 (Zeiss) and imaged using ZEN v3.1 blue edition (Zeiss).

### Quantification of histology and immunostaining

Complete mid-ventricular heart images were quantified using RGB color thresholds in NIS-Elements AR (v5.11, Nikon). Positive staining area was normalized to tissue area, defined by total pixels – white background pixels. Specific RGB thresholds are listed in the Supplementary Materials Major Resources Tables. Trichrome staining was additionally subdivided into regional comparisons. Perivascular regions of interest (ROIs) were defined by tracing the lumens of 2-4 left ventricular free wall coronary arterial branches, dilating each ROI 50 times, then removing the lumen from the ROI using the draw holes function. Perivascular ROIs were then averaged within each mouse prior to statistical analysis. Equivalent-sized rectangular ROIs (860 µm x 430 µm) were drawn in regions corresponding to the anterior, free wall, and posterior epicardium, right ventricle, and anterior, mid, and posterior interventricular septum. Rectangular ROI placement and rotation were adjusted within distinct anatomical regions to maximize inclusion of fibrosis when present.

### Bulk RNA sequencing

Mice were euthanized by a cocktail of ketamine and xylazine (90 mg/kg #11695–6840-1, 10 mg/kg #11695–4024-1, respectively; Covetrus). The right atrial appendage was excised, cold saline (~10 ml) was perfused via the left ventricle, and hearts were harvested immediately. Mid-ventricular sections approximately 1 mm in width were excised using a cold dissection block, rapidly imaged using a Nikon SMZ25 stereoscope (Nikon), and snap-frozen in liquid nitrogen. RNA was extracted using Maxwell RSC simplyRNA Tissue Kits (#AS1340, Promega) according to the manufacturer’s protocol. Samples were processed individually for bulk RNA sequencing, with single mice representing independent experimental replicates. RNA samples were sequenced by Novogene. The sequencing library was generated from total mRNA (1 µg) using NEBNext Ultra™ RNA Library Prep Kits for Illumina (New England BioLabs). cDNA libraries were then sequenced by a Next Generation NovaSeq platform, (HWI-ST1276, Illumina, USA), in a pair-end fashion to reach more than 1,500,000 reads. FASTQ sequence data were mapped to the reference mouse genome using HISAT2 (v2.2.1) and quantified using featureCounts (v2.0.6).

Bulk RNA sequencing data were analyzed on R (v4.5.1) ^27^. Ensembl gene identifiers with no detected read counts in any sample were excluded. Read count data were normalized using the TMM method in the edgeR package (v4.6.3) ^28^ to adjust for biases in library size and composition. Multi-group comparisons were made using “glmLRT” with hemorrhage presence as a binary covariate in edgeR ^28^. Since the present study aimed to profile genes with a high potential for forming pathology in AngII-infused mice, miscellaneous transcripts such as duplicated, unnamed, ribosomal, and mitochondrial genes were removed. Criteria for removal were based on searches within Ensembl gene identifiers for “NA,” duplicated identifiers, or identifiers containing: “Gm[0 – 9],” “[0 – 9]Rik,” “RP[2 – 9],” or “mt-.” Gene set enrichment analysis (GSEA) of mouse ortholog Hallmark gene sets was performed using the fgsea package (v1.34.2) on ranked genes ^29^. Ranks were computed as the sign of the log fold change multiplied by the square root of the likelihood ratio from the interaction model, with sets filtered to 10-500 genes. Mouse ortholog Hallmark gene sets were retrieved from MSigDB with the msigdbr package (v25.1.1) ^30^. The Benjamini–Hochberg method was used for multiple testing correction and an adjusted *p*<0.05 was considered statistically significant. Chord plot illustration was performed using the SRplot online tool ^31^.

## Statistical analysis

Data are presented in scatter plots with down triangles or diamonds representing biological replicates. For parametric data, the mean and standard error of the mean are presented as adjacent circles and error bars. For non-parametric data, the median and 25th/75th percentiles are presented in boxes and 5^th^/95^th^ percentiles in whiskers, with adjacent dots representing outliers. Statistical analyses for all measurements excepting bulk RNA sequencing were performed using SigmaPlot (v16.0, SYSTAT Software). Normality and homogeneity of variance were assessed by Shapiro–Wilk and Brown–Forsythe tests, respectively. Log, inverse, or Box–Cox transformations were applied to data failing normality and Shapiro–Wilk and Brown–Forsythe tests were repeated. For independent measurements with confirmed normal distribution before or after transformation: homoscedastic and heteroscedastic data were analyzed by Student’s and Welch’s *t* tests, respectively. Data failing tests for normality and homogeneity of variance were analyzed by Mann–Whitney U test. For repeated measurements, all data met assumptions for normality and homogeneity of variance either before or after transformation and were analyzed by two-way repeated measures ANOVA. *p*≤0.05 was considered statistically significant.

### Sex as a biological variable

For 1 and 4-week infusion pathological studies using PAI-1+/+ and PAI-1−/− mice, both male and female animals were examined, and similar findings were reported for both sexes. Since no pronounced sexual dimorphism was observed in AngII or NE-induced pathology, only male mice were examined for the 1-week bulk RNA sequencing and 1-day AngII infusion studies or for studies in PAI-1^Ala/Ala^ and PAI-1^AK/AK^ mice.

### Study approval

All experiments were performed according to a protocol approved by the Institutional Animal Care and Use Committee at the University of Kentucky (Protocol #2018-2967) in accordance with the guidelines of the National Institutes of Health.

### Data availability

RNA sequencing data (raw FASTQ and aligned data) will be made publicly available at the gene expression omnibus (GEO) repository. Numerical data that support the findings of this study are provided in the Supporting Data Values file. The data that support the findings of this study are also available from the corresponding authors on reasonable request.

## RESULTS

### PAI-1 deficiency failed to alter ascending thoracic aortic aneurysm (ATAA) in mice

AngII infusion induces ascending aortic dilatation in mice, recapitulating multiple pathological features of ATAA ^32^. Our previous work demonstrated that PAI-1 protein abundance was increased markedly in the ascending aortas of mice infused with AngII for 3 days, a stage before overt pathology ^33^. Of clinical relevance, PAI-1 was also highly abundant in human ATAA tissue ^33^. To determine whether PAI-1 contributes to ATAA, AngII was infused for 4 weeks in both male and female PAI-1−/− and their PAI-1+/+ littermates starting at 8-14 weeks of age (Figure 1A). Despite the increased abundance of PAI-1 in thoracic aortas during aneurysm formation ^33^, its absence did not alter AngII-induced ATAA. No difference was detected between genotypes in ascending aortic diameters, as measured both in live animals by ultrasound (Figure 1B) and *in situ* at termination (Figure 1C). To assess whether PAI-1 deficiency influenced AngII-induced aortic wall remodeling, ascending and descending thoracic aortas were stained for collagen and elastin. Collagen deposition and elastin fragmentation were not changed between PAI-1+/+ and −/− mice infused with AngII (Figure 1, D and E).

**Figure 1.**
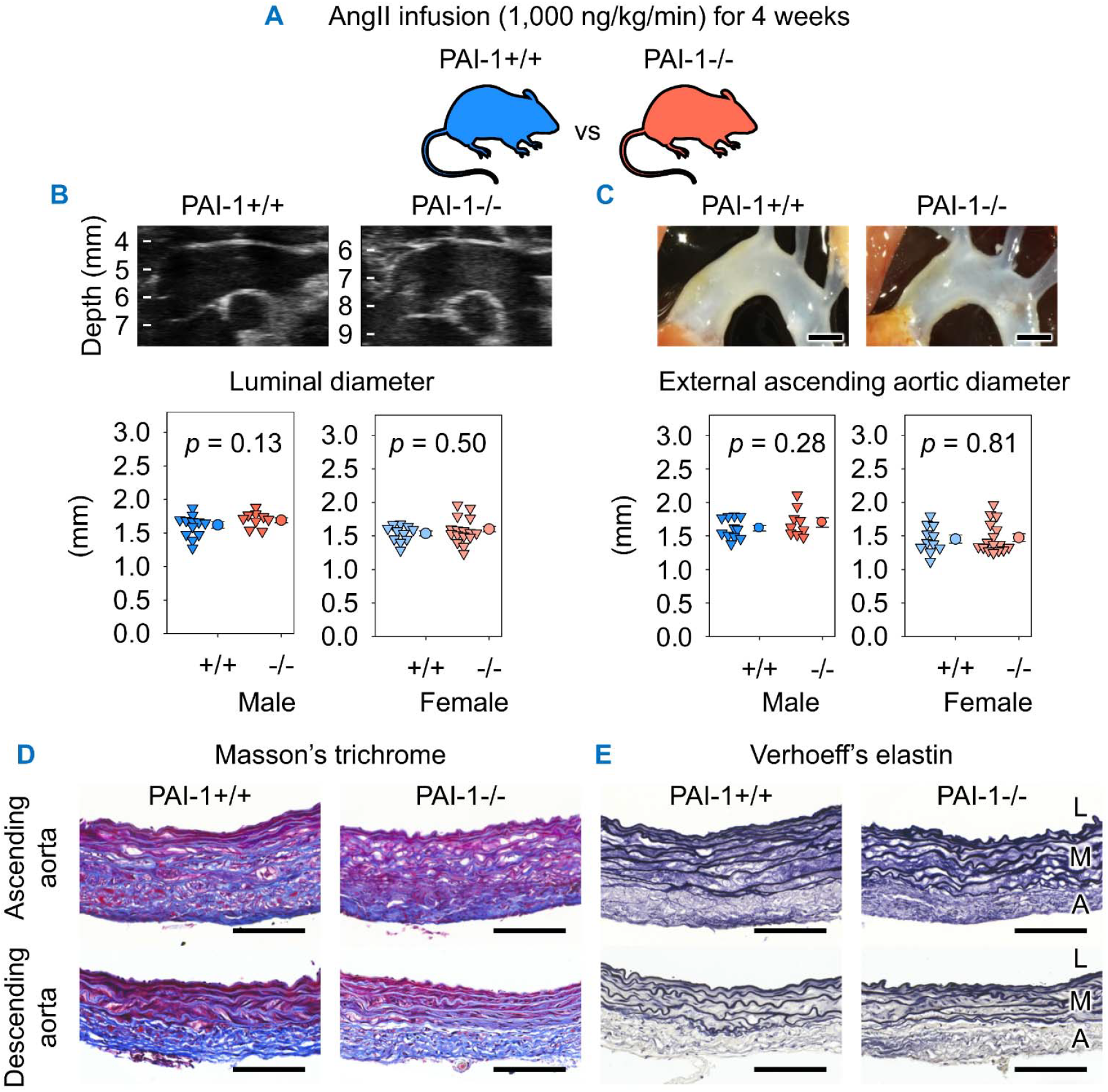
PAI-1 deficiency failed to alter ascending thoracic aortic aneurysm in mice. (**A**) Male and female PAI-1+/+ and −/− littermates were infused with AngII (1,000 ng/kg/min) for 4 weeks. (**B**) Representative right parasternal ultrasound images and quantification of maximal luminal ascending aortic diameter measurements. (**C**) Representative *in situ* aortic images and quantification of maximal external aortic diameter measurements. Representative Masson’s trichrome (**D**) and Verhoeff’s elastin staining (**E**) of ascending and descending aortic sections. Representative images are from male mice. Comparisons between genotypes made by Student’s *t* test (**B**) and Student’s *t* test for males and Student’s *t* test after log transformation for females (**C**). *n* = 9-16 mice/group. Scale bars = 1,000 µm and 100 µm for whole section and high-magnification images, respectively. Abbreviations: AngII: angiotensin II, PAI-1: plasminogen activator inhibitor-1, L: lumen, M: media, A: adventitia.

### PAI-1 deficiency augmented AngII-induced cardiac fibrosis and led to ferric iron deposition

While investigating the contribution of PAI-1 to ATAA, grossly evident cardiac remodeling was observed in both male and female AngII-infused PAI-1−/− mice (Figure 2, A and B). Histological examination of mid-ventricular sections demonstrated increased cardiac collagen deposition in PAI-1−/− mice in patterns consistent with both interstitial and replacement fibrosis (Figure 2C). To assess the distribution of fibrosis, select cardiac regions of interest (ROIs) were compared between genotypes (Supplemental Figure 1A and B). Collagen was localized primarily to the epicardium of both ventricles and the posterior septum (Supplemental Figure 1C-F, I). Myocardial fibrosis was also present consistently but distributed heterogeneously. Male PAI-1−/− mice displayed a modest increase in fibrosis in the anterior and mid septum (Supplemental Figure 1G-H). In contrast, perivascular fibrosis was not altered by PAI-1 deficiency (Supplemental Figure 1J).

**Figure 2.**
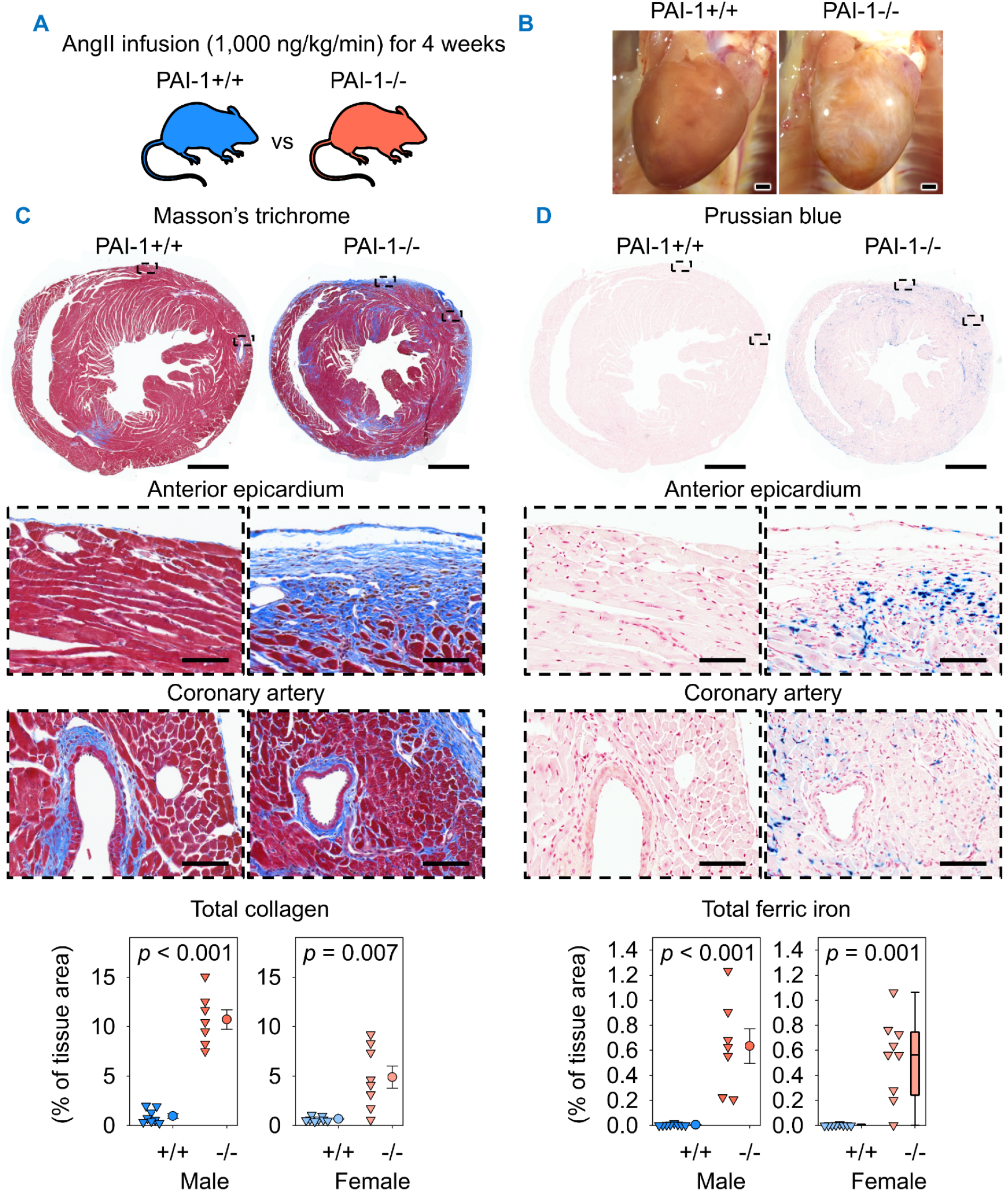
PAI-1 deficiency augmented AngII-induced cardiac fibrosis and induced ferric iron deposition. (**A**) Male and female PAI-1+/+ and −/− littermates were infused with AngII (1,000 ng/kg/min) for 4 weeks. (**B**) Representative *in situ* heart images. (**C**) Representative Masson’s trichrome staining and collagen quantification of mid-ventricular hearts. Comparisons between genotypes were made by Welch’s *t* test. (**D**) Representative Prussian blue staining and ferric iron quantification of mid-ventricular hearts. Comparisons between genotypes were made by Student’s *t* test after log transformation for males and Mann-Whitney U test for females. *n* = 7-9 mice/group. Representative images are from male mice. Scale bars = 1,000 µm and 100 µm for whole section and high-magnification images, respectively. Abbreviations: AngII: angiotensin II, PAI-1: plasminogen activator inhibitor-1.

Given the role of PAI-1 in hemostasis, ferric iron deposition was examined as a marker of prior hemorrhage. PAI-1−/− mice exhibited ferric iron deposition in regions spatially associated with collagen accumulation, particularly within the epicardium. In contrast, ferric iron accumulation was minimal in PAI-1+/+ mice. Quantification of Prussian blue staining confirmed that PAI-1 deficiency markedly increased ferric iron deposition (Figure 2D). Combined picrosirius red, fast green, and Prussian blue staining demonstrated colocalization of ferric iron with both interstitial and replacement fibrosis (Supplemental Figure 2).

To investigate whether PAI-1 deficiency augments fibrosis or promotes ferric iron deposition in other highly vascularized organs, tissue sections were examined from the lung, liver, and kidney. Previous studies have reported that PAI-1 deficiency reduces fibrosis in these organs under pathological conditions ^6^. No differences were detected in collagen deposition between PAI-1+/+ and −/− mice (Supplemental Figure 3, A-C). Similarly, no apparent ferric iron deposition was detected in either genotype within serial tissue sections of lung, liver, and kidney (Supplemental Figure 4, A-C). Together, these findings indicate that PAI-1 deficiency promotes AngII-induced ferric iron deposition and exacerbates fibrosis in the heart but not in other organs.

To determine whether PAI-1 deficiency alone was sufficient to induce cardiac pathology, 8 to 14-week-old male and female mice were infused with saline (the vehicle of AngII) for 4 weeks (Supplemental Figure 5A). Cardiac remodeling, determined by gross examination and Masson’s trichrome staining, was not evident in PAI-1−/− mice (Supplemental Figure 5, B and C). Furthermore, cardiac ferric iron deposition was not evident in either genotype (Supplemental Figure 5D). These data demonstrate that PAI-1 deficiency does not induce cardiac pathology in young mice under basal conditions, but markedly augments cardiac fibrosis and induces ferric iron deposition in response to AngII infusion.

### PAI-1 deficiency did not affect systolic function during AngII infusion

The extent of hypertension-induced cardiac remodeling in PAI-1−/− mice prompted us to investigate whether PAI-1 deficiency also impaired cardiac function. Therefore, male and female mice were infused with either saline or AngII and cardiac morphometry, systolic function, and wall strain were measured by echocardiography at baseline and during 1 and 4 weeks of infusion (Supplemental Figure 6A). No alterations were detected due to the interaction of group and infusion time in left ventricular internal diameter, calculated left ventricular mass, left ventricular anterior wall thickness, ejection fraction, or global longitudinal or circumferential strain. Modest increases in left ventricular posterior wall thickness were detected by interaction analysis in male and female PAI-1−/− mice infused with AngII (Supplemental Figure 6, B and C). Overall, despite the pronounced cardiac pathology in PAI-1−/− mice, no significant differences were discerned in systolic function or wall strain at the intervals studied.

### PAI-1 deficiency induced cardiac fibrosis and ferric iron deposition in norepinephrine-infused mice

Infusion of AngII at a rate of 1,000 ng/kg/min increases systolic blood pressure in mice ^34–36^. To evaluate whether hypertension, per se, promotes cardiac remodeling observed in AngII-infused PAI-1−/− mice, both male and female mice at 11-12 weeks of age were infused with a pressor dose of norepinephrine (NE) for 4 weeks (Figure 3A). Gross cardiac remodeling was evident in PAI-1−/− mice, irrespective of sex, after 4 weeks of NE infusion (Figure 3B). PAI-1+/+ mice exhibited minimal cardiac collagen deposition following NE infusion. In contrast, PAI-1−/− mice displayed interstitial and replacement fibrosis. Quantification of collagen deposition confirmed that PAI-1 deficiency induced marked cardiac fibrosis (Figure 3C). Fibrosis was further characterized in select ROIs (Supplemental Figure 7A and B). PAI-1−/− mice displayed pronounced collagen accumulation in the anterior epicardium and right ventricle in both sexes and in the posterior septum in males (Supplemental Figure 7C, F, and I). Modest increases in collagen were observed in the free wall and posterior epicardium, anterior and mid septum, and perivascular regions of PAI-1−/− mice (Supplemental Figure 7D, E, G, H, and J). Ferric iron deposition was detected in PAI-1−/− mice in spatial association with collagen deposition, and quantification of ferric iron staining confirmed its increased abundance (Figure 3D). Ferric iron was observed coincident with replacement and interstitial fibrosis by combined picrosirius red, fast green, and Prussian blue staining (Supplemental Figure 8).

**Figure 3.**
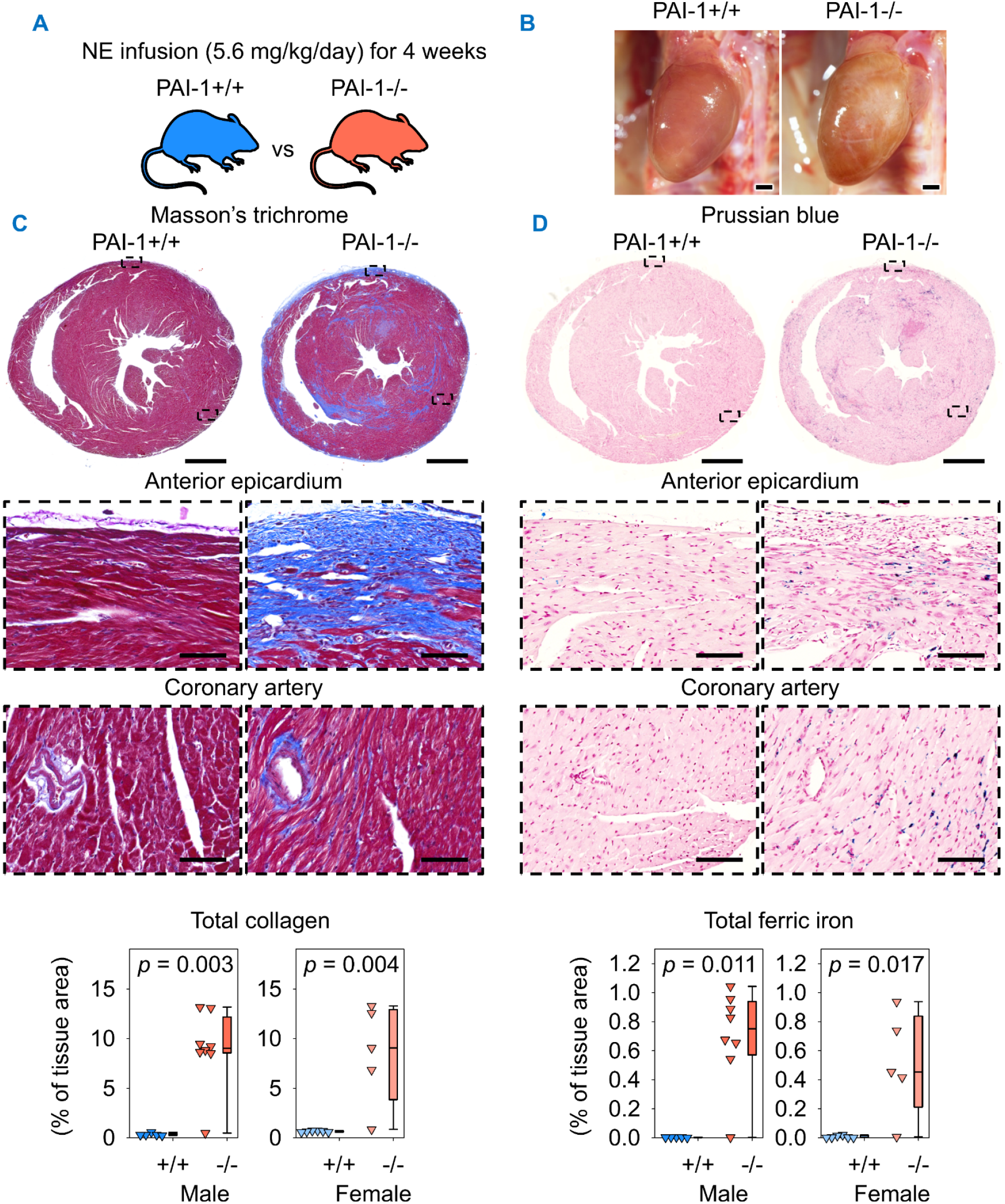
PAI-1 deficiency induced cardiac fibrosis and ferric iron deposition in NE-infused mice. (**A**) Male and female PAI-1+/+ and −/− littermates were infused with NE (5.6 mg/kg/day) for 4 weeks. (**B**) Representative *in situ* heart images. Representative Masson’s trichrome staining and collagen quantification (**C**) and Prussian blue staining and ferric iron quantification (**D**) of mid-ventricular hearts. Comparisons between genotypes were made by Mann-Whitney U test. *n* = 5-8 mice/group. Representative images are from male mice. Scale bars = 1,000 µm and 100 µm for whole section and high-magnification images, respectively. Abbreviations: NE: norepinephrine, PAI-1: plasminogen activator inhibitor-1.

### PAI-1 deficiency induced cardiac hemorrhage and fibrosis after 1 week of AngII or NE infusion

To determine early pathological events associated with cardiac remodeling in PAI-1−/− mice, both male and female PAI-1+/+ and −/− mice at 11-12 weeks of age were infused with AngII for 1 week (Figure 4A), an interval associated with acute aortic pathology ^32^. Grossly evident cardiac hemorrhage was observed in PAI-1−/− mice, whereas hemorrhage was barely detectable in PAI-1+/+ mice (Figure 4B). Immuno-histochemical staining for TER-119, a marker of erythrocytes, demonstrated extensive cardiac hemorrhage in PAI-1−/− mice, most pronounced within the epicardium and also present within the myocardium. Quantification of TER-119-positive area confirmed the increased hemorrhage in PAI-1−/− mice (Figure 4C). This early pathological event was associated spatially with the augmented epicardial fibrosis observed after 4 weeks of AngII infusion.

**Figure 4.**
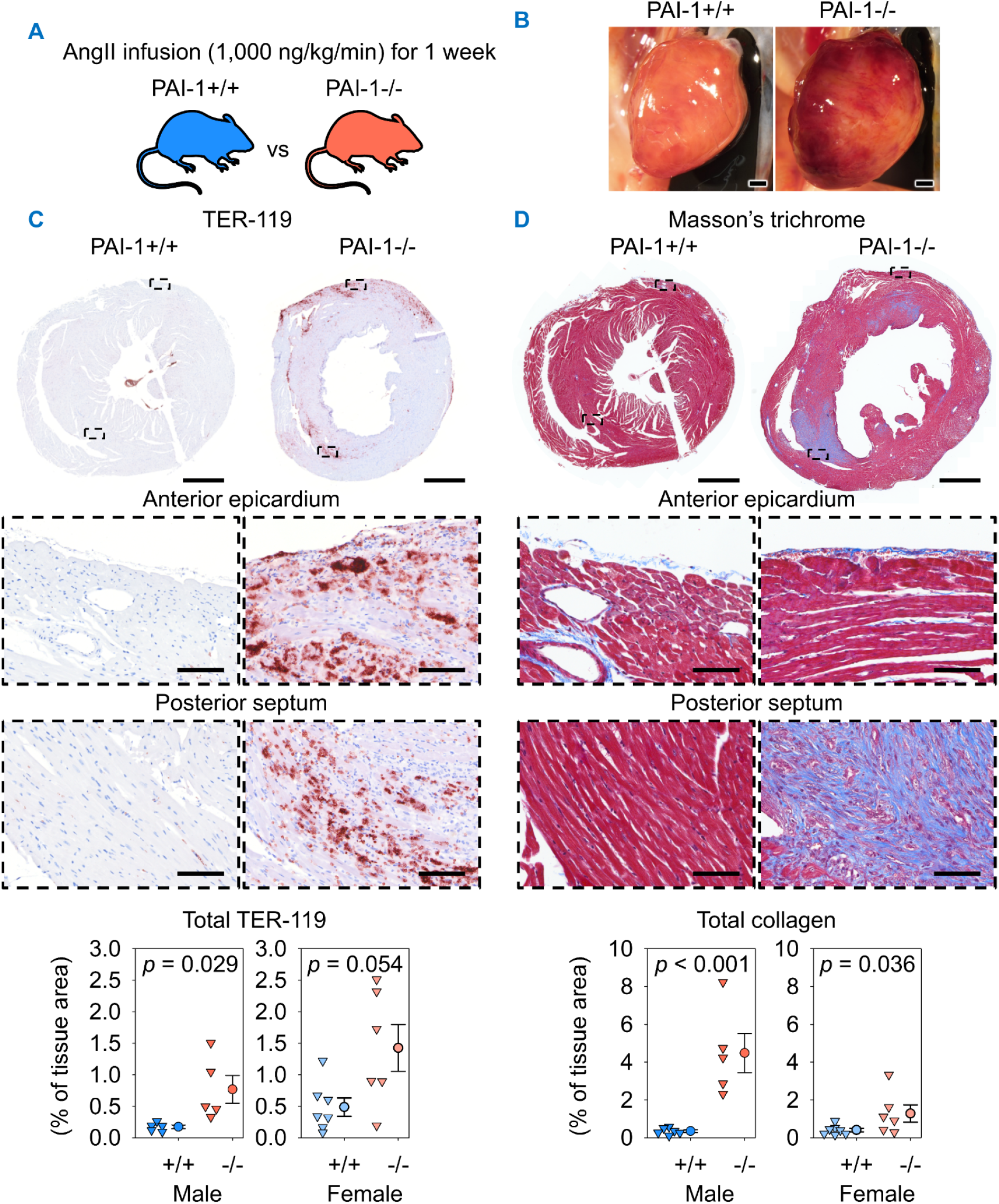
PAI-1 deficiency augmented AngII-induced cardiac fibrosis and promoted hemorrhage after 1 week of AngII infusion. (**A**) Male and female PAI-1+/+ and −/−littermates were infused with AngII (1,000 ng/kg/min) for 1 week. (**B**) Representative *in situ* heart images. (**C**) Representative TER-119 immunohistochemistry and quantification of mid-ventricular hearts. Comparisons between genotypes were made by Student’s *t* test for males and Welch’s *t* test for females. (**D**) Masson’s trichrome staining and collagen quantification of mid-ventricular hearts. Comparisons between genotypes were made by Student’s *t* test after log transformation. *n* = 5-7 mice/group. Representative images are from male mice. Scale bars = 1,000 µm and 100 µm for whole section and high-magnification images, respectively. Abbreviations: AngII: angiotensin II, PAI-1: plasminogen activator inhibitor-1.

One week of AngII infusion also resulted in significant cardiac fibrosis in PAI-1−/− mice. Collagen deposition was observed predominantly in the myocardium, particularly within the posterior septum. Quantification of Masson’s trichrome staining confirmed augmented cardiac fibrosis in PAI-1−/− mice (Figure 4D). Hemorrhage was observed coincident with fibrotic remodeling in 3 of 5 PAI-1−/−males and 4 of 6 PAI-1−/−females. Ferric iron deposition was also increased in PAI-1−/− mice in association with fibrotic regions such as the posterior septum and adjacent to hemorrhagic regions within the epicardium (Supplemental Figure 9).

The above findings were also confirmed in both male and female mice infused with NE for 1 week (Figure 5A). NE-infused PAI-1−/− mice developed grossly evident cardiac hemorrhage, while no visible hemorrhage was detected in PAI-1+/+ mice (Figure 5B). The distribution of hemorrhage, as detected by TER-119 staining, was predominantly within the epicardium and myocardium, consistent with the distribution in PAI-1−/− mice infused with AngII for 1 week. Quantification of the TER-119-positive area verified the increased hemorrhage in PAI-1−/− mice (Figure 5C). Masson’s trichrome staining revealed that collagen deposition in PAI-1−/− mice was localized mainly to the myocardium, including the posterior septum, a distribution similar to mice infused with AngII for 1 week. Quantification of collagen-positive area demonstrated increased fibrosis in PAI-1−/− mice after 1 week of NE infusion (Figure 5D). Hemorrhage was observed coincident with fibrosis in 5 of 6 male and 5 of 6 female PAI-1−/− mice. Ferric iron was also increased in PAI-1−/− mice, localized both in fibrotic and hemorrhagic regions within the posterior septum and epicardium, respectively (Supplemental Figure 10).

**Figure 5.**
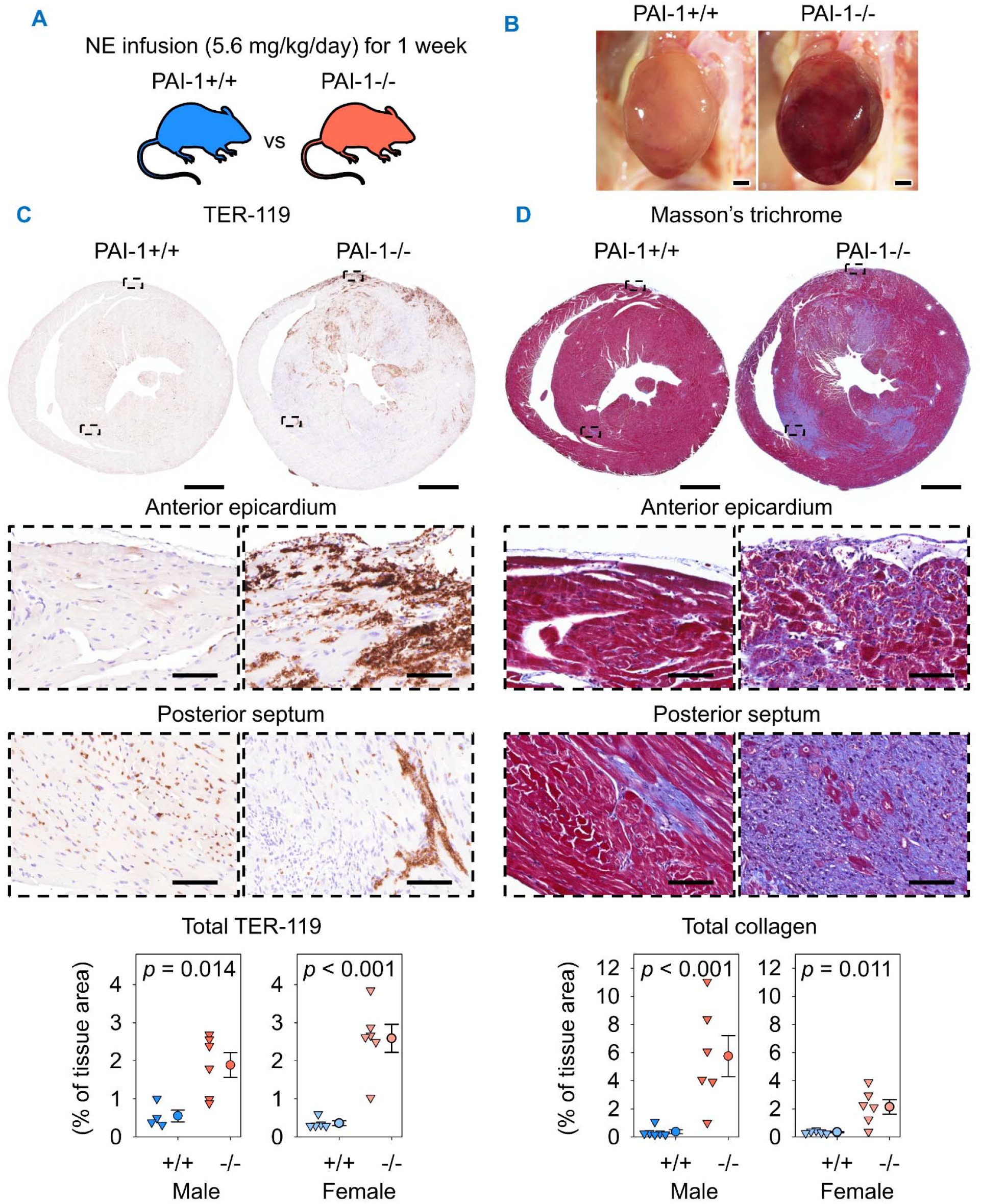
PAI-1 deficiency augmented NE-induced cardiac fibrosis and promoted hemorrhage after 1 week of NE infusion. (**A**) Male and female PAI-1+/+ and −/−littermates were infused with NE (5.6 mg/kg/day) for 1 week. (**B**) Representative *in situ* heart images. (**C**) Representative TER-119 immunohistochemistry and quantification of mid-ventricular hearts. Comparisons between genotypes were made by Student’s *t* test for males and Student’s *t* test after log transformation for females. (**D**) Representative Masson’s trichrome staining and collagen quantification of mid-ventricular hearts. Comparisons between genotypes were made by Student’s *t* test after log transformation for males and Student’s *t* test for females. *n* = 5-6 mice/group. Representative images are from male mice. Scale bars = 1,000 µm and 100 µm for whole section and high-magnification images, respectively. Abbreviations: NE: norepinephrine, PAI-1: plasminogen activator inhibitor-1.

### PAI-1 deficiency increased transcripts related to proteolysis and fibrosis independently of hemorrhage severity after 1 week of AngII infusion

To determine the transcriptional alterations associated with cardiac pathology in PAI-1−/− mice, male PAI-1+/+ and −/− mice were infused with saline or AngII for 1 week. Since both male and female PAI-1−/− mice displayed increased cardiac hemorrhage and fibrosis in response to hypertension, only male mice were used. Mid-ventricular tissue sections were analyzed by bulk RNA-sequencing. Among PAI-1−/− mice infused with AngII, 3 mice displayed modest hemorrhage within the sequenced sample, while 7 mice showed pronounced hemorrhage (Supplemental Figure 11A). To avoid complicated data interpretation due to ranges of hemorrhage severity, the presence of prominent hemorrhage was used as a binary covariate in the RNA sequencing analysis, which detected 34,196 genes.

Principal component analysis using unfiltered transcriptomes revealed distinct alterations in AngII-infused mice compared to saline infusion. Within AngII-infused mice, PAI-1−/− mice displayed further transcriptomic alteration from baseline compared to PAI-1+/+ mice (Supplemental Figure 11B). Interaction analysis between infusion and genotype detected 82 differentially expressed genes (DEGs). Hierarchical clustering identified 5 distinct gene clusters (Supplemental Figure 11C). DEGs within cluster #1 were downregulated the most within AngII-infused PAI-1−/− mice. Cluster #2 was comprised of genes upregulated within AngII-infused PAI-1+/+ mice. DEGs within cluster #3 displayed increased expression coincident with significant hemorrhage presence in AngII-infused PAI-1−/− mice. Cluster #4 contained a single gene, *C1qtnf3*, that was highly expressed within AngII-infused PAI-1−/− mice with modest hemorrhage (Supplemental Figure 11D). The largest number of DEGs were grouped within cluster #5; these genes were highly abundant in AngII-infused PAI-1−/− mice, irrespective of hemorrhage severity.

To investigate the potential contribution of plasmin to the observed pathology, we next performed targeted analysis for proteolytic substrates or regulatory transcripts. Interestingly, *Fn1* was the only plasmin(ogen)-related transcript within the 82 interaction DEGs and was grouped within cluster #5. Therefore, we next analyzed our dataset for genes differentially expressed due to genotype or infusion. In addition to *Serpine1* (PAI-1), only *C1qtnf3* was downregulated in PAI-1−/− mice, indicating that PAI-1 deficiency minimally alters the cardiac transcriptome at baseline. Analysis for transcripts altered by AngII infusion revealed 2,603 DEGs. Among these, 21 DEGs related to plasmin, which were subdivided into four categories: positive (Supplemental Figure 12A) or negative regulation of plasminogen activation (Supplemental Figure 12B), proteolytic targets likely to promote extracellular matrix turnover (Supplemental Figure 12C), and proteolytic targets likely to induce intracellular signaling (Supplemental Figure 12D).

To identify individual transcripts with a high potential for forming pathology in AngII-infused PAI-1−/− mice, the differential expression analysis was integrated with gene set enrichment analysis focused on mouse ortholog Hallmark gene sets. Among the top 10 Hallmark gene sets with positive enrichment scores, 12 DEGs were present within the leading edge of the gene sets (Supplemental Figure 13A). All 12 DEGs were present within cluster #5. Additionally, 6 of the 12 DEGs were present within at least two Hallmark gene sets and encoded for ECM components (*Fn1* and *Ecm1*), proteases (*Lgals3* and *Ctss*), a ferroptosis-associated polyamine catabolic enzyme (*Sat1*), or an immune-stimulatory receptor (*Cd48*, Supplemental Figure 13B). These findings suggest that PAI-1 deficient hearts exhibit transcriptional signatures of injury and remodeling in response to AngII, independent of the severity of hemorrhage. Furthermore, these results support that hypertension-induced cardiac injury may be initiated prior to 1 week of infusion.

### PAI-1 deficiency augmented cardiac hemorrhage and myocyte injury within 1 day of AngII infusion

PAI-1 deficiency promoted fibrosis, iron accumulation, and hemorrhage after 1 week of AngII or NE infusion. To determine whether hemorrhage occurs rapidly, 9 to 12-week-old mice were infused with AngII for 1 day (Figure 6A). Since both male and female PAI-1−/− mice displayed increased cardiac hemorrhage and fibrosis in response to AngII infusion for 1 and 4 weeks, only male mice were infused for the 1-day interval. Grossly evident cardiac hemorrhage was detected in the posterior wall of PAI-1−/− mice, while hemorrhage was barely visible in PAI-1+/+ mice (Figure 6B). Staining for erythrocytes revealed diffuse hemorrhage localized within the myocardium and posterior septum of PAI-1−/− mice (Figure 6C). This distribution of hemorrhage was consistent with the distribution of collagen accumulation in PAI-1−/− mice after 1 week of AngII infusion. Quantification of TER-119-positive area confirmed the augmentation of hemorrhage in PAI-1−/− mice (Figure 6C). Masson’s trichrome staining and quantification confirmed that hemorrhage occurred prior to collagen accumulation in PAI-1−/− mice (Figure 6D).

**Figure 6.**
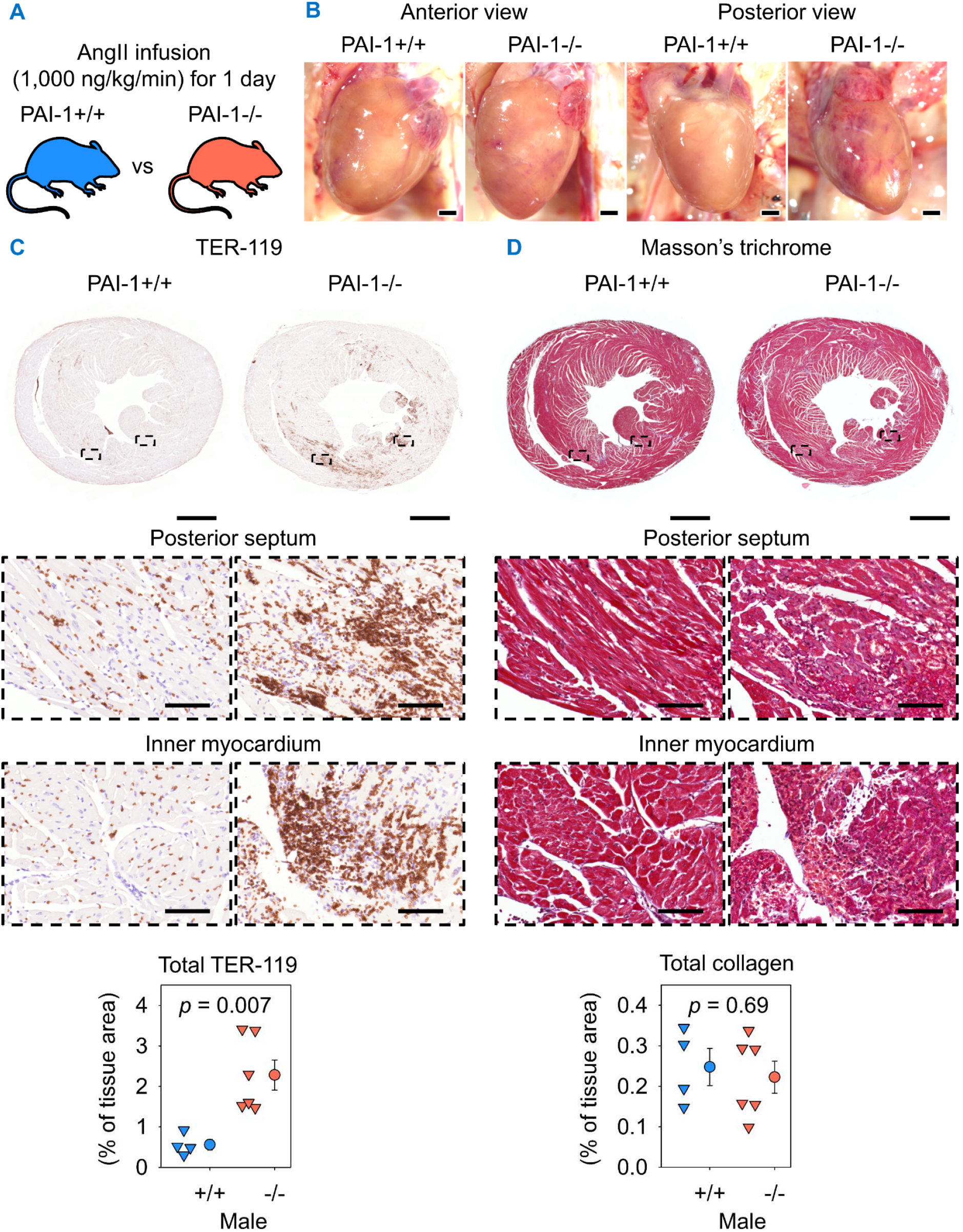
PAI-1 deficiency induced cardiac hemorrhage after 1 day of AngII infusion. (**A**) Male PAI-1+/+ and −/− littermates were infused with AngII (1,000 ng/kg/min) for 1 day. (**B**) Representative *in situ* heart images. (**C**) Representative TER-119 immunostaining and quantification in mid-ventricular heart sections. (**D**) Representative Masson’s trichrome staining and collagen quantification of mid-ventricular hearts. Comparisons were made by Student’s *t* test. *n* = 4-6 mice/group. Scale bars = 1,000 µm and 100 µm for whole section and high-magnification images, respectively. Abbreviations: AngII: angiotensin II, PAI-1: plasminogen activator inhibitor-1.

The replacement fibrosis observed after 7 and 28 days of AngII or NE-induced blood pressure increase led us to investigate whether cardiomyocyte injury was evident at 1 day of AngII infusion. Therefore, staining for cardiac troponin I, a marker of cardiomyocytes, was performed. Restricted areas of nucleated, non-vascular, troponin I-negative myocardium were observed in PAI-1+/+ mice, consistent with minor cardiomyocyte injury (Supplemental Figure 14). In contrast, cardiomyocyte injury was evident prominently in PAI-1−/− mice, particularly within the posterior septum and myocardium. Serial staining of TER-119 and troponin I indicated that areas of cardiomyocyte injury in PAI-1−/− mice coincided predominantly with hemorrhage (Figure 6C and Supplemental Figure 14). To verify that TER-119 accumulation in PAI-1−/− mice was representative of hemorrhage, co-immunofluorescent (co-IF) staining was performed for nuclei (4’,6-diamidino-2-phenylindole, DAPI), endothelium (CD31), and erythrocytes (TER-119). Erythrocytes within the myocardium of PAI-1−/− mice accumulated outside the vessel lumens, indicating hemorrhage (Supplemental Figure 15, A and B).

### Mutational disruption of the RCL, but not the SMB-binding domain, of PAI-1 contributed to AngII-induced cardiac injury

The RCL of PAI-1 is responsible for its protease inhibitory function ^4, 20, 37^. In contrast, the distal SMB-binding domain of PAI-1 mediates its interaction with vitronectin ^18, 19^. To examine the role of these structural domains in hemorrhage and fibrosis, male mice harboring loss-of-function point mutations of either domain were infused with saline or AngII for 1 week (Supplemental Figure 16A and Figure 7A). Mice homozygous for mutations in the SMB-binding domain of PAI-1 (R101A and Q123K; PAI-1^AK/AK^) did not display gross cardiac hemorrhage or TER-119 accumulation after AngII infusion (Supplemental Figure 16B and C). Additionally, collagen deposition was not increased in AngII-infused PAI-1^AK/AK^ mice (Supplemental Figure 16D). Ferric iron deposition was minimal in PAI-1^AK/AK^ mice, suggesting an absence of prior hemorrhage (Supplemental Figure 17). These results indicate that the SMB-binding domain of PAI-1 is not required for its protective effect.

**Figure 7.**
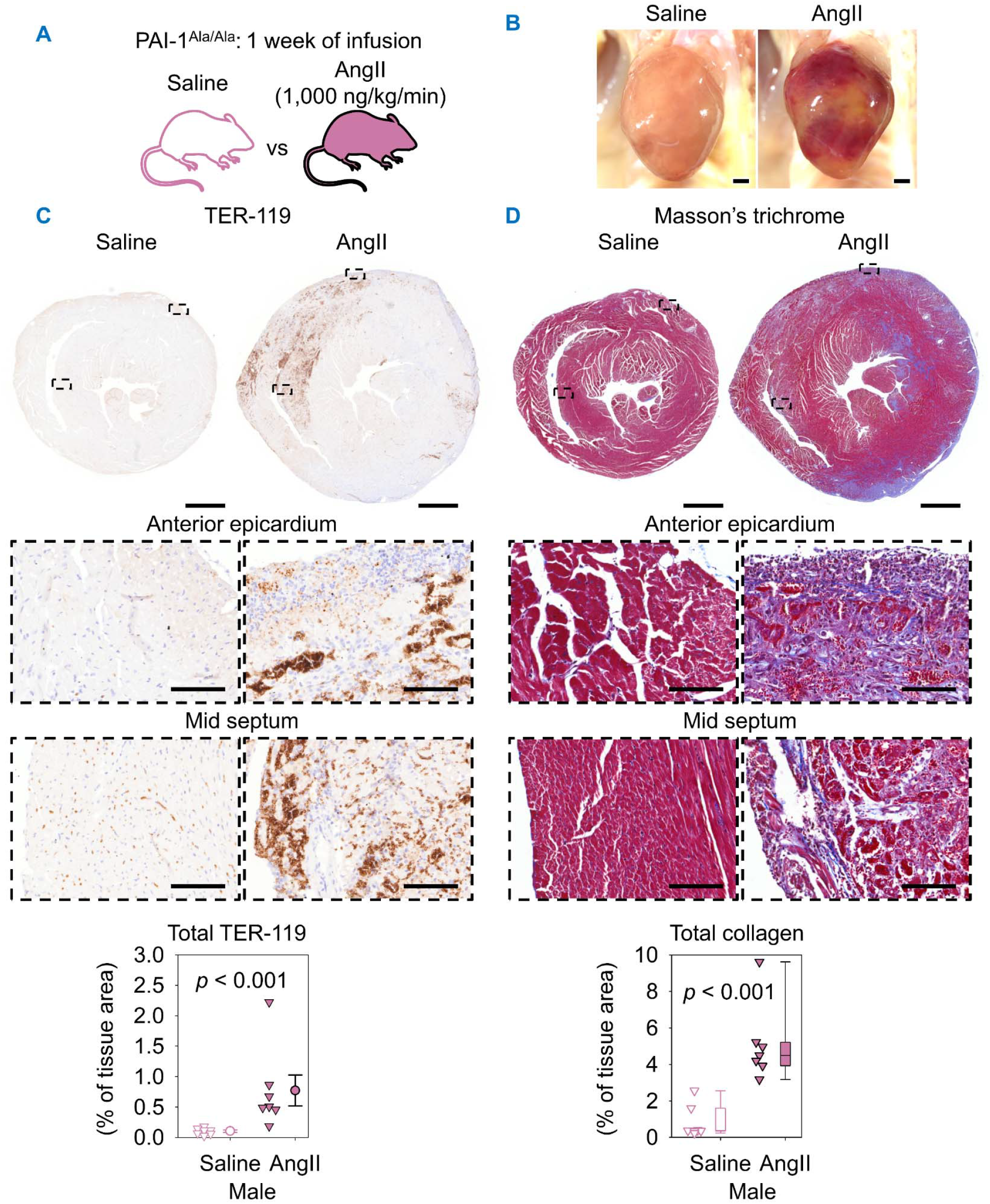
Mutational disruption of the reactive center loop domain of PAI-1 contributed to AngII-induced cardiac injury. (**A**) Male PAI-1Ala/Ala littermates were infused with saline or AngII (1,000 ng/kg/min) for 1 week. (**B**) Example *in situ* heart images. (**C**) Example TER-119 immunohistochemistry and quantification of mid-ventricular hearts. Comparison was made by Student’s *t* test after log transformation. (**D**) Masson’s trichrome staining and collagen quantification of mid-ventricular hearts. Comparison was made by Mann-Whitney U test. *n* = 7 mice/group. Scale bars = 1,000 µm and 100 µm for whole section and high-magnification images, respectively. Abbreviations: AngII: angiotensin II, PAI-1: plasminogen activator inhibitor-1.

In contrast, mice homozygous for a mutation in the RCL (R346A; PAI-1^Ala/Ala^) displayed grossly evident cardiac hemorrhage after 1 week of AngII (Figure 7B). Erythrocyte accumulation, as detected by TER-119 immunostaining, was increased by AngII infusion and localized primarily to the epicardium and mid septum (Figure 7C). AngII also increased collagen deposition in PAI-1^Ala/Ala^ mice, with a heterogeneous distribution among mice; prominent collagen deposition was detected in the septum, myocardium, and epicardium (Figure 7D). Hemorrhage was observed coincident with fibrosis in 6 of 7 AngII-infused PAI-1^Ala/Ala^ mice. Ferric iron deposition was also induced by AngII infusion and localized to hemorrhagic and fibrotic areas (Supplemental Figure 18). These findings support the conclusion that the protease inhibitory function of PAI-1 protects against AngII-induced cardiac injury.

## DISCUSSION

The initial objective of this study was to assess the role of PAI-1 in AngII-induced ATAA. PAI-1 deficiency, however, did not affect ascending aortic dilatation or vascular pathologies after 4 weeks of AngII infusion. Instead, cardiac fibrosis was observed in PAI-1−/− mice in response to AngII infusion. While this finding has been reported previously ^8, 11–13^, our studies, with detailed spatial and temporal exploration, revealed several novel observations: (1) PAI-1 deficiency augmented cardiac fibrosis and promoted ferric iron deposition in both AngII- and NE-infused mice, regardless of sex; (2) PAI-1 deficiency triggered cardiac hemorrhage and cardiomyocyte injury as early as 1 day of AngII infusion; (3) hemorrhage observed at 1 day and 1 week infusion intervals spatially coincided with augmented fibrosis observed at 1 and 4 week intervals, respectively; and (4) disruption of the RCL, but not the SMB-binding domain, of PAI-1 induced cardiac hemorrhage and fibrosis at 1 week of AngII infusion.

Despite the abundance of PAI-1 in human ATAA and ascending aortas of AngII-infused mice ^33^, its deficiency did not alter AngII-induced aortic dilatation or remodeling. This outcome differs from reports of reduced aortic remodeling in AngII-infused PAI-1−/− mice ^8, 13^. In addition to the aorta, PAI-1 deficiency augmented coronary perivascular fibrosis after 4 weeks of NE, but not AngII, infusion in our study, differing from the protective effect of PAI-1 deficiency against Nω-Nitro-L-arginine methyl ester (L-NAME)-induced perivascular fibrosis ^38, 39^. These discrepancies likely reflect the distinct vascular stresses and stimuli induced by different models of hypertension ^40^. Additional variables such as age at study onset, AngII infusion strategy, and strain differences may influence the impact of PAI-1 deficiency on vascular remodeling.

In the heart, extensive fibrosis was observed in PAI-1−/− mice after 1 or 4 weeks of AngII or NE infusion, consistent with previous reports in AngII-infused mice ^8, 11–13^. Our studies exhibited that PAI-1 deficiency augmented cardiac fibrosis regardless of sex, whereas analysis stratified by sex has not been reported previously. In the present study, 8 to 14-week-old mice were used, which exhibited no detectable cardiac fibrosis in the absence of AngII or NE infusion, replicating previous work in mice within 6 months of age ^9, 13^. However, spontaneous cardiac fibrosis has been reported in 9 to 24-month-old PAI-1−/− mice ^7, 9, 10^. These data reveal a remarkable contrast: PAI-1 deficiency alone induces gradual cardiac fibrosis that is only manifest at advanced age, whereas infusions with either AngII or NE generate dramatic pathology within 1 to 4 weeks. To determine the associated functional consequences, we performed longitudinal echocardiography during AngII infusion. PAI-1−/− mice had increased posterior wall thickness, consistent with previously reported hypertrophy ^13^. Systolic function was unchanged by PAI-1 deficiency, contrasting with the reduced ejection fraction reported after 2 weeks of AngII followed by 2 weeks without infusion in 16 to 20-week-old mice ^13^. Differences in age at study onset and AngII infusion intervals may influence the susceptibility of PAI-1−/− mice to AngII-induced systolic dysfunction. Overall, our findings demonstrate that hypertension induced cardiac fibrosis and modest alterations in ventricular wall thickness without detectable impairment of systolic function.

Either in aged ^7, 9, 10^ or AngII-infused mice ^8, 11–13^, PAI-1 deficiency does not induce fibrosis in the lung, liver, or kidney. Therefore, the acceleration of fibrosis appears to be unique to the hearts of PAI-1−/− mice. The heart is a highly muscular organ that constantly contracts to propel blood to the circulatory system, exposing it to substantial mechanical load. Given that both AngII and NE, two molecules well known to increase blood pressure, exacerbate cardiac fibrosis in PAI-1−/− mice, a plausible accelerating mechanism is constant hemodynamic mechanical stress. Such stress may promote microinjury and amplify fibrotic signaling cascades in cardiac tissue lacking PAI-1, ultimately precipitating the pronounced fibrotic remodeling observed in these mice.

To further understand the potential mechanisms by which PAI-1 accelerates AngII-augmented cardiac fibrosis in mice, AngII was infused for either 1 or 7 days. These investigations in early infusion intervals of either AngII or NE build upon prior reports of cardiac fibrosis in PAI-1 deficiency ^7–13^ by identifying hemorrhage and myocyte injury as preceding events. PAI-1−/− mice displayed diffuse posterior septal and myocardial hemorrhage and myocyte injury after only 1 day of AngII infusion. After 1 week of AngII or NE infusion, PAI-1−/− mice displayed primarily replacement fibrosis within the myocardium and posterior septum as well as diffuse epicardial hemorrhage. Ferric iron accumulation was evident coincident with fibrosis in the septum and adjacent to hemorrhage in the epicardium. Transcriptomic profiling after 1 week of AngII revealed elevations in fibrotic and proteolytic transcripts independent of hemorrhage severity, consistent with pathway-level results reported previously ^13^. By 4 weeks of AngII or NE infusion, PAI-1−/− mice displayed predominant epicardial replacement fibrosis, in addition to myocardial and posterior septal fibrosis. Ferric iron deposition was coincident with interstitial and replacement fibrosis, resembling findings in aged mice or young adult mice co-administered with AngII and aldosterone ^9, 12^. Of note, this phenotype has not been reported previously in mice receiving infusions of either NE or AngII. The predominantly epicardial distribution of injury resembles that seen in conditional prothrombin deficient mice ^41^, aged low-tissue-factor or low-factor VII mice ^42, 43^, and in cardiomyocyte-specific tissue factor-deficient mice exposed to isoproterenol ^44^. Cardiac iron accumulation and fibrosis in PAI-1−/− mice is also comparable in magnitude to the predominantly myocardial pathology observed in aged factor X Friuli mice ^45^. Such regional injury and our observations of diffuse hemorrhage likely reflect microvascular susceptibility to hemodynamic stress. AngII and NE infusion produced similar pathologies in PAI-1−/− mice after 1 and 4 weeks of infusion, indicating that hemodynamic stress itself, rather than hormone-specific signaling, drives the phenotype, paralleling a report of increased cardiac fibrosis in cardiomyocyte-specific PAI-1−/− mice after transverse aortic constriction ^46^. These results highlight the unique vulnerability of the heart to stress-induced microvascular damage. Observations of hemorrhage being spatially coincident with subsequent fibrosis implicate microvascular instability as a driver of cardiac injury in PAI-1−/− mice.

PAI-1 is a multifunctional serine protease inhibitor that reduces plasmin-mediated fibrinolysis and extracellular proteolysis through inhibition of plasminogen activators via its RCL ^4, 20, 37^. PAI-1 also exerts several plasmin-independent effects as a regulator of cell adhesion, migration, and angiogenesis via domains distinct from its RCL ^47, 48^. In contrast to its effect in the heart, PAI-1 often promotes organ fibrosis ^6^; PAI-1 mutational studies have demonstrated that it promotes lung fibrosis independently of its RCL ^18, 19^. Prior reports have implicated uPA and plasmin-mediated activation of transforming growth factor-β as key drivers of cardiac microvascular injury and fibrosis in PAI-1 deficiency ^12, 13^. Our studies in PAI-1^Ala/Ala^ mice demonstrate that disruption of the RCL induces cardiac hemorrhage, iron deposition, and fibrosis after 1 week of AngII, with regional injury similar to that observed after 1 week of AngII or NE infusion in PAI-1−/− mice. These findings align with the reduction of AngII+aldosterone-induced cardiac iron accumulation and fibrosis in double deficient uPA−/−& PAI-1−/− or Pg^S743A/S743A^ & PAI-1−/− mice ^12, 13^. In contrast, PAI-1^AK/AK^ mice did not have detectable cardiac injury after 1 week of AngII, supporting that the SMB-binding domain, and resultant stabilization of PAI-1 in its active conformation by vitronectin, is not required for its protective effect. These results differ from a previous report in which recombinant PAI-1 AK protein augmented cardiac fibrosis in uninephrectomized C57BL/6 mice fed a high-salt diet and infused with AngII ^49^. These divergent results may be explained by the absence or presence of native PAI-1, similar to the observed concentration-dependent pro- or anti-angiogenic effects of PAI-1 ^50, 51^. Collectively, our data and previous studies support a model in which PAI-1 deficiency results in dysregulated plasmin activity that promotes cardiac microvascular injury and subsequent fibrosis.

In summary, this study demonstrated that PAI-1 deficiency promoted hypertension-induced cardiac hemorrhage and fibrosis triggered by two distinct stimuli. Spatial, temporal, and molecular analyses support a model in which microvascular hemorrhage and cardiomyocyte injury precede fibrotic remodeling in the absence of PAI-1 or disruption of its RCL. These findings underscore a critical role for PAI-1 in preserving cardiac microvascular integrity under hemodynamic stress and highlight the protective function of the RCL. In agreement with these experimental studies, cardiac fibrosis has also been reported in a human population with PAI-1 deficiency ^13, 52^, highlighting the clinical relevance of the phenotype observed in mice. Cardiac fibrosis resulting from PAI-1 deficiency currently lacks targeted therapy. Our findings suggest that these patients may be intrinsically susceptible to hemodynamic stress-induced cardiac microvascular injury. Approaches aimed at minimizing elevations in blood pressure, promoting endothelial stability, or reducing plasmin activity could therefore represent additional therapeutic strategies.

## AUTHOR CONTRIBUTIONS

Conceptualization: ACP, DAL, THS, HS, JES, HSL, AD

Data curation: ACP, SI, HS, HSL, AD

Formal analysis: ACP, DBG, VZG, NZ, HS, JES

Funding acquisition: ACP, AD

Investigation: ACP, SI, MKF, DAH, JJM, BML, DBG, VZG, NZ, JES

Methodology: ACP, HS, HSL, AD

Project administration: HSL, AD

Writing–original draft: ACP

Writing–review & editing: SI, MKF, DAH, JJM, BML, DBG, VZG, NZ, DAL, THS, HS, JES, HSL, AD

## ACKNOWLEDGMENTS

None

