## Supplemental Materials for "Cardiac Hemorrhage Precedes Hypertension-induced Fibrosis in Plasminogen Activator Inhibitor-1 Deficient Mice"

Hong S. Lu

Saha CVRC, BBSRB, Room 249

University of Kentucky

741 S Limestone

Lexington KY 40536-0509

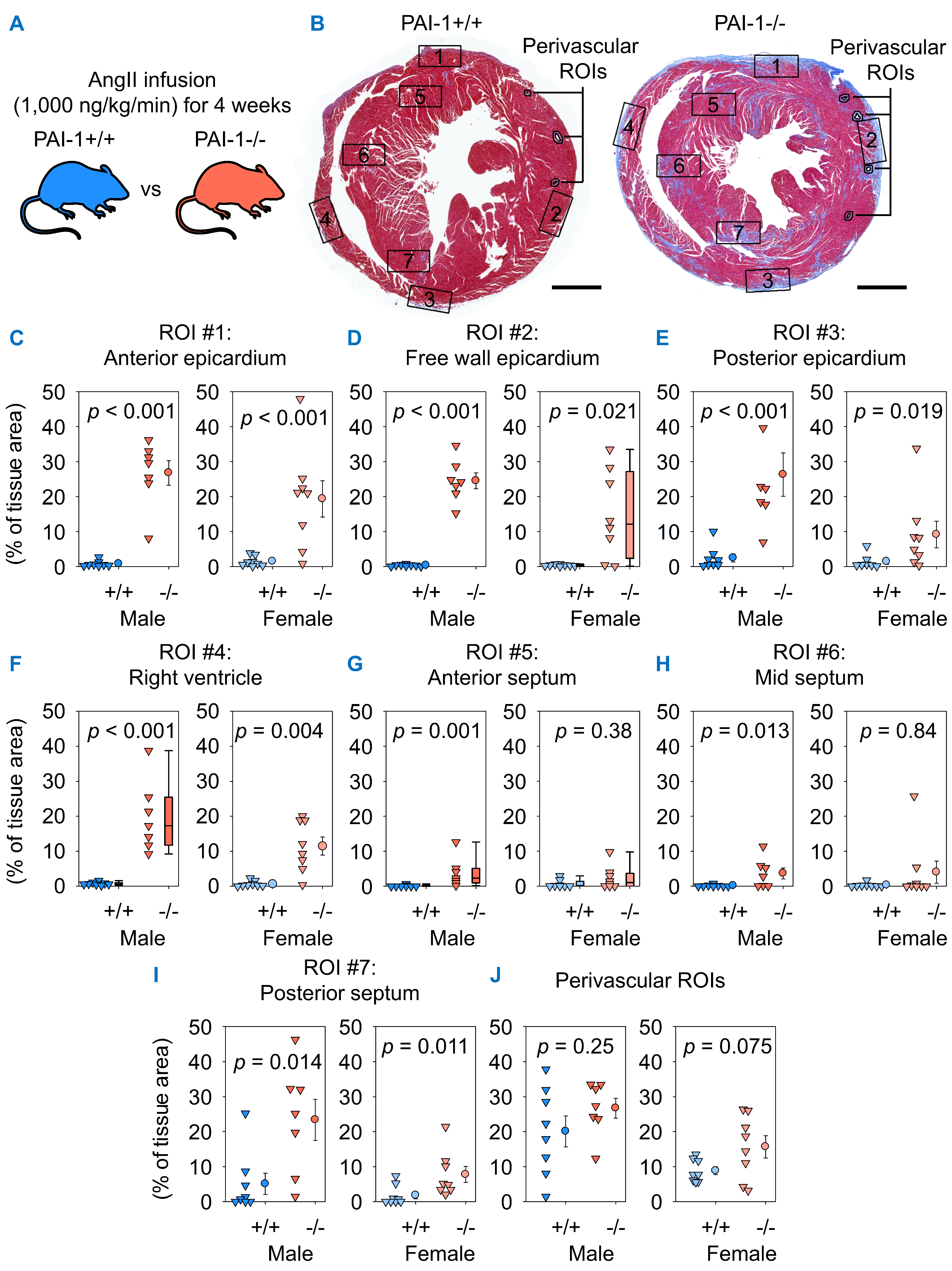

**Supplemental Figure 1.** **PAI-1 deficiency augmented epicardial and septal fibrosis after 4 weeks of AngII infusion.** (**A**) Male and female PAI-1+/+ and -/- littermates were infused with AngII for 4 weeks. (**B**) Representative ROI placement on Masson’s trichrome-stained mid-ventricular heart sections. Collagen quantification within ROI #1: anterior epicardium (**C**), ROI #2: free wall epicardium (**D**), ROI #3: posterior epicardium (**E**), ROI #4: right ventricle (**F**), ROI #5: anterior septum (**G**), ROI #6: mid septum (**H**), ROI #7: posterior septum (**I**), and perivascular ROIs (**J**). Comparisons between genotypes were made by: Student’s *t* test before (**I-J**) or after log-transformation (**C-E, H**) or Mann-Whitney U test (**F-G**) for males and Student’s *t* test after log transformation (**C, E**), Welch’s *t* test before (**F, J**) or after log transformation (**H-I**), or Mann-Whitney U test (**D, G**) for females. *n* = 7-9 mice/group. Scale bars = 1,000 µm. Abbreviations: AngII: angiotensin II, PAI-1: plasminogen activator inhibitor-1, ROI: region of interest.

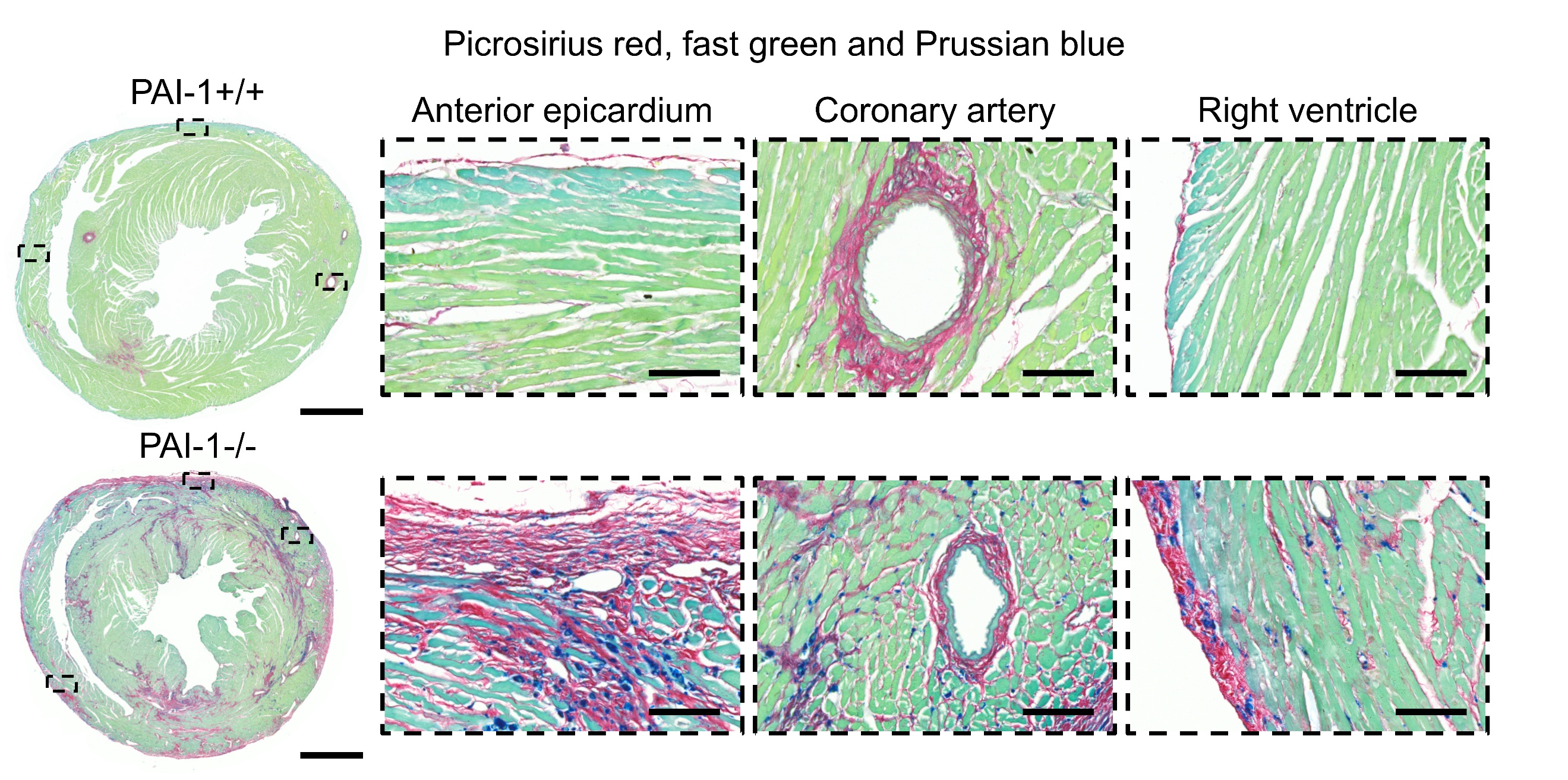

**Supplemental Figure 2.** **Ferric iron deposits were associated with fibrosis in PAI-1 deficient mice after 4 weeks of AngII infusion.** Representative picrosirius red, fast green, and Prussian blue co-staining of mice infused with AngII (1,000 ng/kg/min) for 4 weeks. Representative images are from male mice. Scale bars = 1,000 µm and 100 µm for whole section and high-magnification images, respectively. Abbreviations: AngII: angiotensin II, PAI-1: plasminogen activator inhibitor-1.

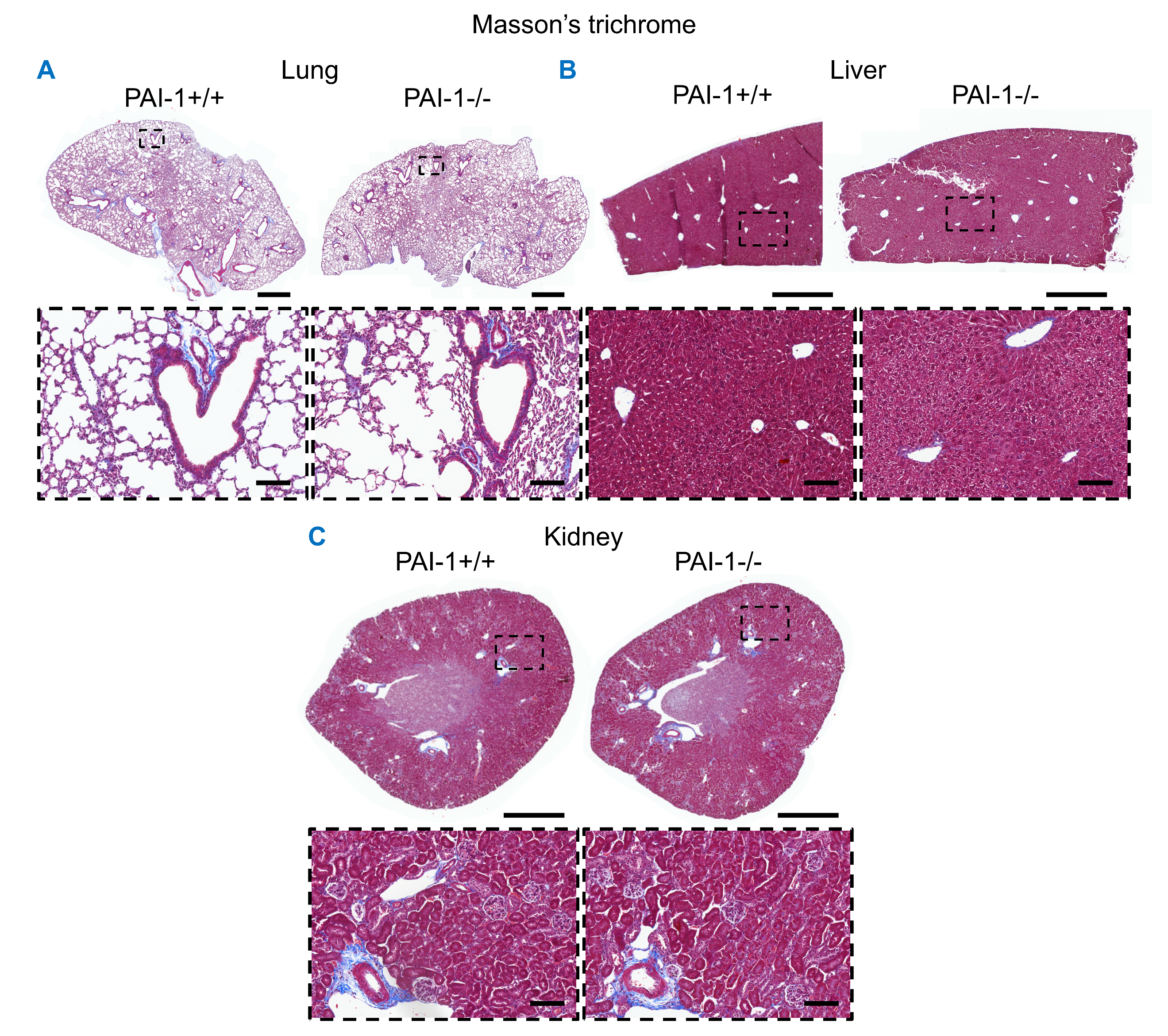

**Supplemental Figure 3.** **PAI-1 deficiency did not overtly alert lung, liver, or kidney fibrosis.** (**A-C**) Representative Masson’s trichrome staining of left lung (**A**), left liver lobe (**B**), and left kidney sections (**C**) of mice infused with AngII (1,000 ng/kg/min) for 4 weeks. Representative images are from male mice. Scale bars = 1,000 µm and 100 µm for whole section and high-magnification images, respectively. Abbreviations: AngII: angiotensin II, PAI-1: plasminogen activator inhibitor-1.

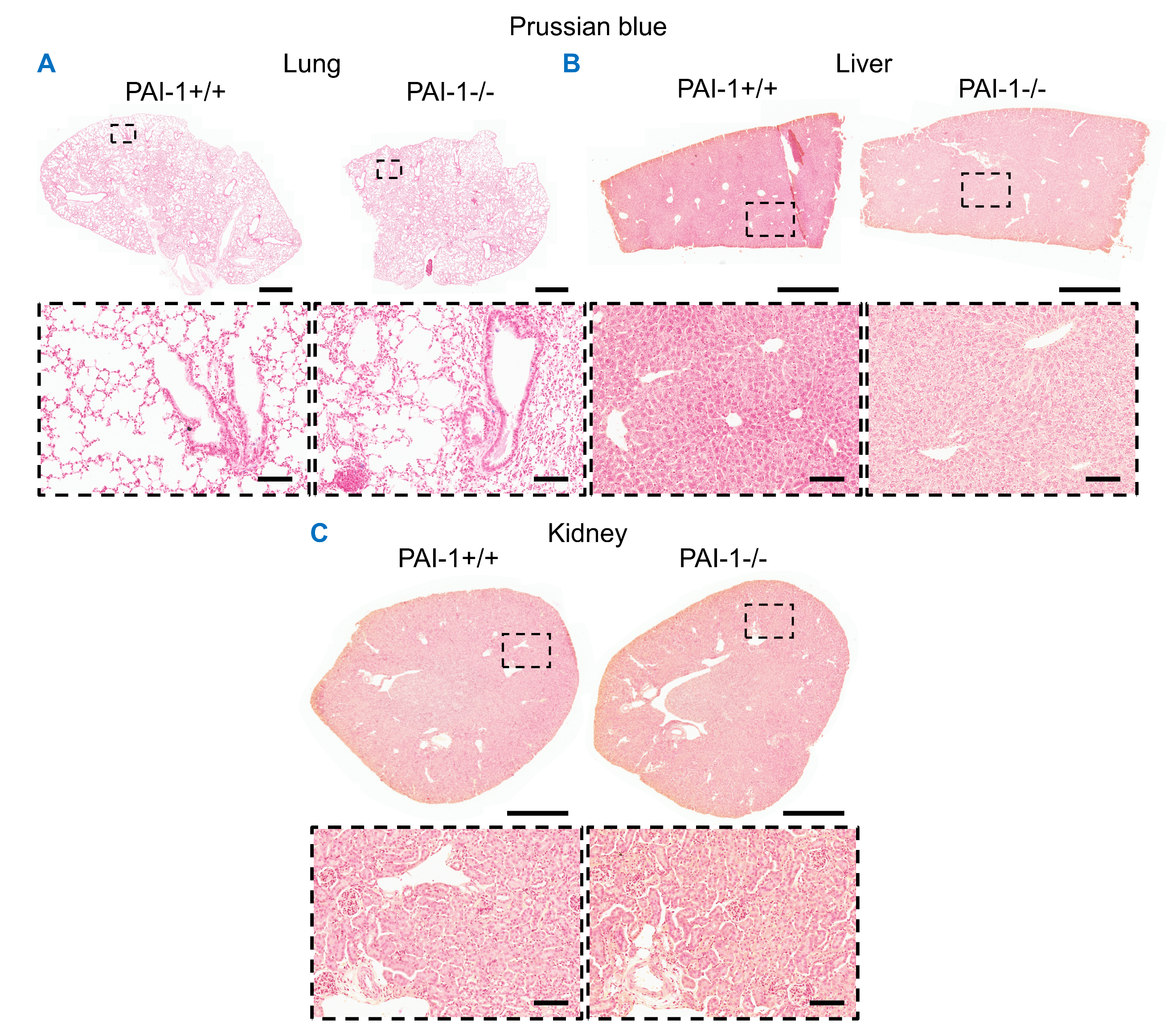

**Supplemental Figure 4.** **PAI-1 deficiency did not detectably alert lung, liver, or kidney ferric iron deposition**. (**A-C**) Representative Prussian blue staining of left lung (**A**), left liver lobe (**B**), and left kidney sections (**C**) of mice infused with AngII (1,000 ng/kg/min) for 4 weeks. Representative images are from male mice. Scale bars = 1,000 µm and 100 µm for whole section and high-magnification images, respectively. Abbreviations: AngII: angiotensin II, PAI-1: plasminogen activator inhibitor-1.

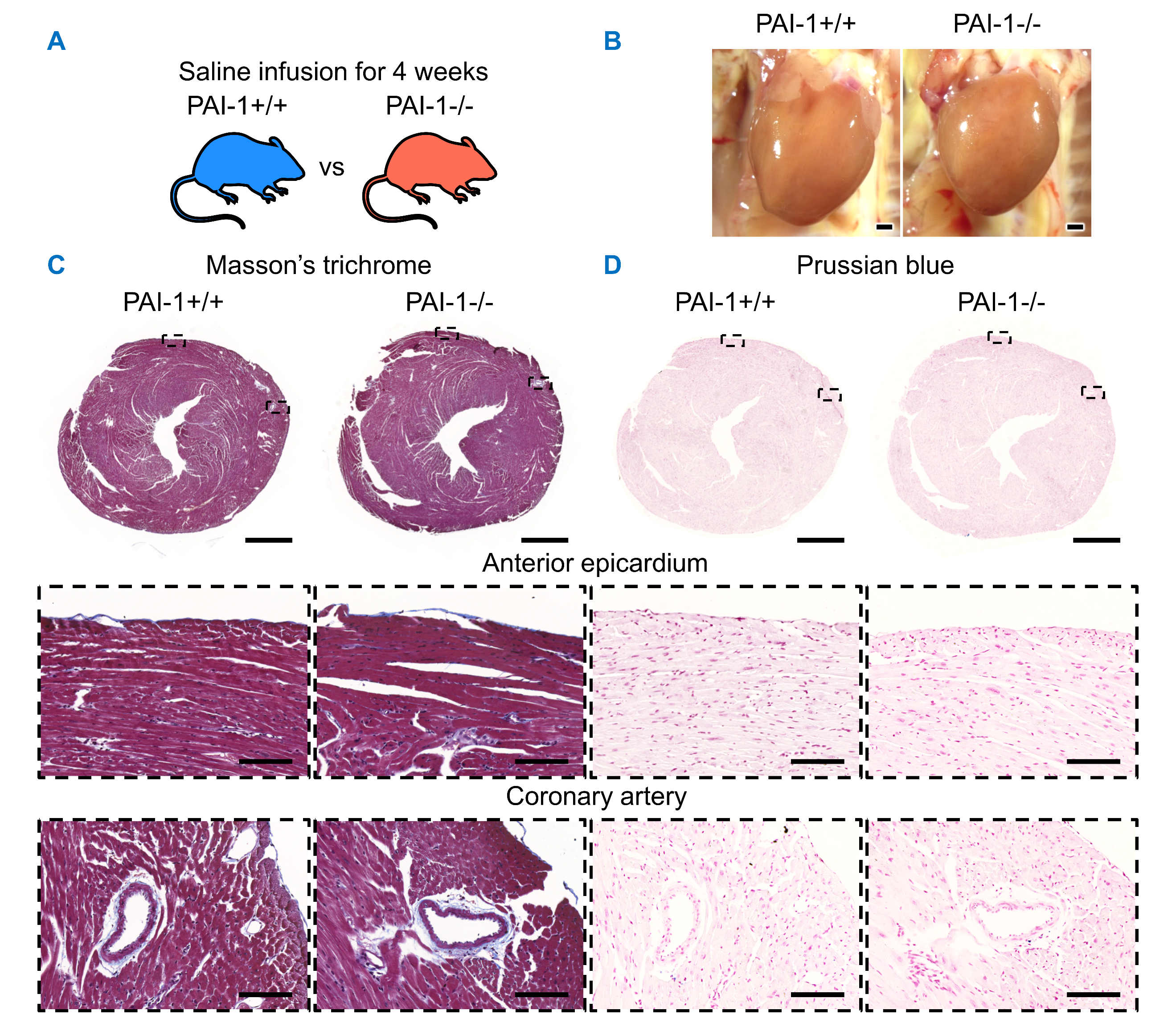

**Supplemental Figure 5**. **PAI-1 deficiency caused minimal cardiac pathology after 4 weeks of saline infusion.** (**A**) Male and female PAI-1+/+ and -/- littermates were infused with saline for 4 weeks. (**B**) Representative *in situ* heart images. (**C-D**) Representative Masson’s trichrome (**C**) and Prussian blue staining (**D**) of hearts. Representative images are from male mice. Scale bars = 1,000 µm and 100 µm for whole section and high-magnification images, respectively. Abbreviations: PAI-1: plasminogen activator inhibitor-1.

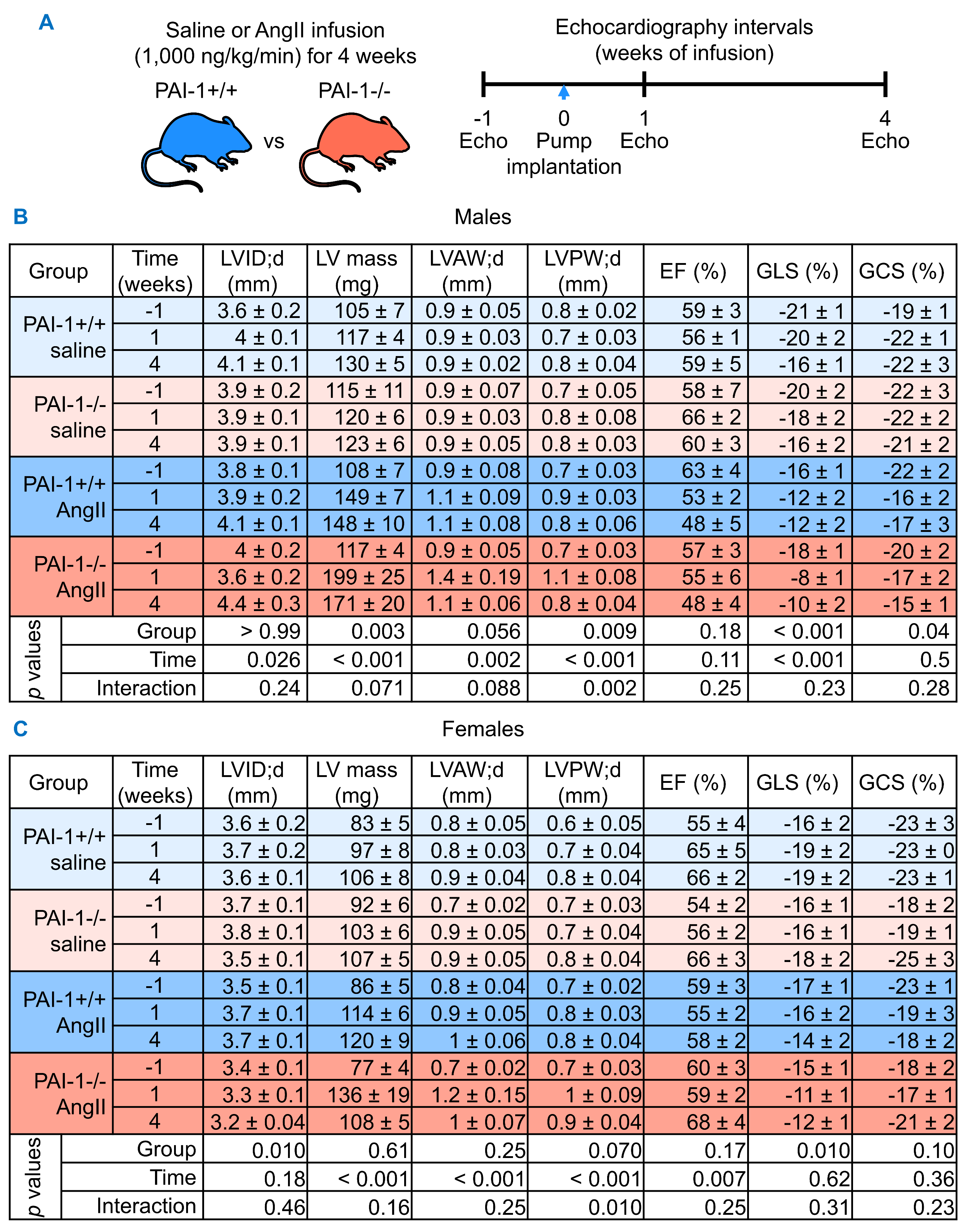

**Supplemental Figure 6.** **PAI-1 deficiency did not alter systolic function within 4 weeks of AngII infusion.** (**A**) Male PAI-1+/+ and -/- littermates were infused with saline or AngII (1,000 ng/kg/min) for 4 weeks. Echocardiography was performed 1 week prior to pump implant (-1) and at 1 and 4 weeks of infusion. (**B-C**) Summary tables of echocardiography parameters in male (**B**) and female mice (**C**). The following transformations were applied prior to analysis: for males: LVPW;d, EF, GLS, and GCS: no transformation; LVID;d: inverse; LV mass and LVAW;d: Box–Cox; for females: LVID;d, GLS, and GCS: no transformation; LV mass: inverse; LVAW;d, LVPW;d, and EF: Box–Cox. Comparisons between groups were made by two-way repeated measures ANOVA. (**B**) *n* = 7 – 8 mice/group. (**C**) *n* = 4 – 10 mice/group. Abbreviations: AngII: angiotensin II, PAI-1: plasminogen activator inhibitor-1, Echo: echocardiography, LVID;d: left ventricular internal diameter; diastole, LV: left ventricle, LVAW;d: left ventricle anterior wall; diastole, LVPW;d left ventricle posterior wall; diastole, EF: ejection fraction, GLS: global longitudinal strain, GCS: global circumferential strain.

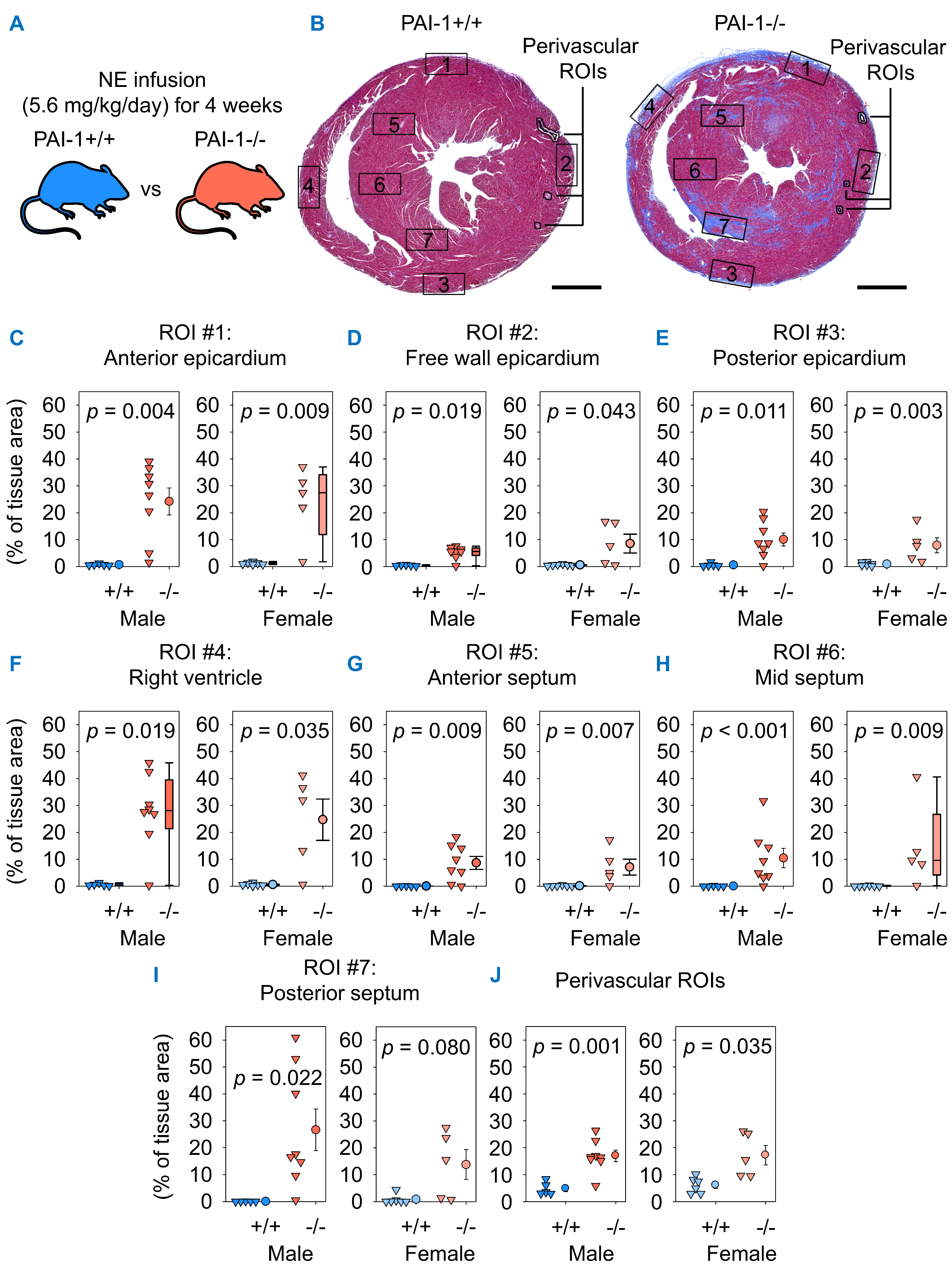

**Supplemental Figure 7.** **PAI-1 deficiency augmented epicardial, septal, and perivascular fibrosis after 4 weeks of NE infusion.** (**A**) Male and female PAI-1+/+ and -/- littermates were infused with NE for 4 weeks. (**B**) Representative ROI placement on Masson’s trichrome-stained mid-ventricular heart sections. Collagen quantification within ROI #1: anterior epicardium (**C**), ROI #2: free wall epicardium (**D**), ROI #3: posterior epicardium (**E**), ROI #4: right ventricle (**F**), ROI #5: anterior septum (**G**), ROI #6: mid septum (**H**), ROI #7: posterior septum (**I**), and perivascular ROIs (**J**). Comparisons between genotypes were made by: Student’s *t* test before (**C, E, I-J**) or after log-transformation (**H**), Welch’s *t* test (**G**), or Mann-Whitney U test (**D, F**) for males and Student’s *t* test after log transformation (**E, G**), Welch’s *t* test before (**F, I-J**) or after log transformation (**D**), and Mann-Whitney U test (**C, H**) for females. *n* = 5-8 mice/group. Scale bars = 1,000 µm. Abbreviations: NE: norepinephrine, PAI-1: plasminogen activator inhibitor-1, ROI: region of interest.

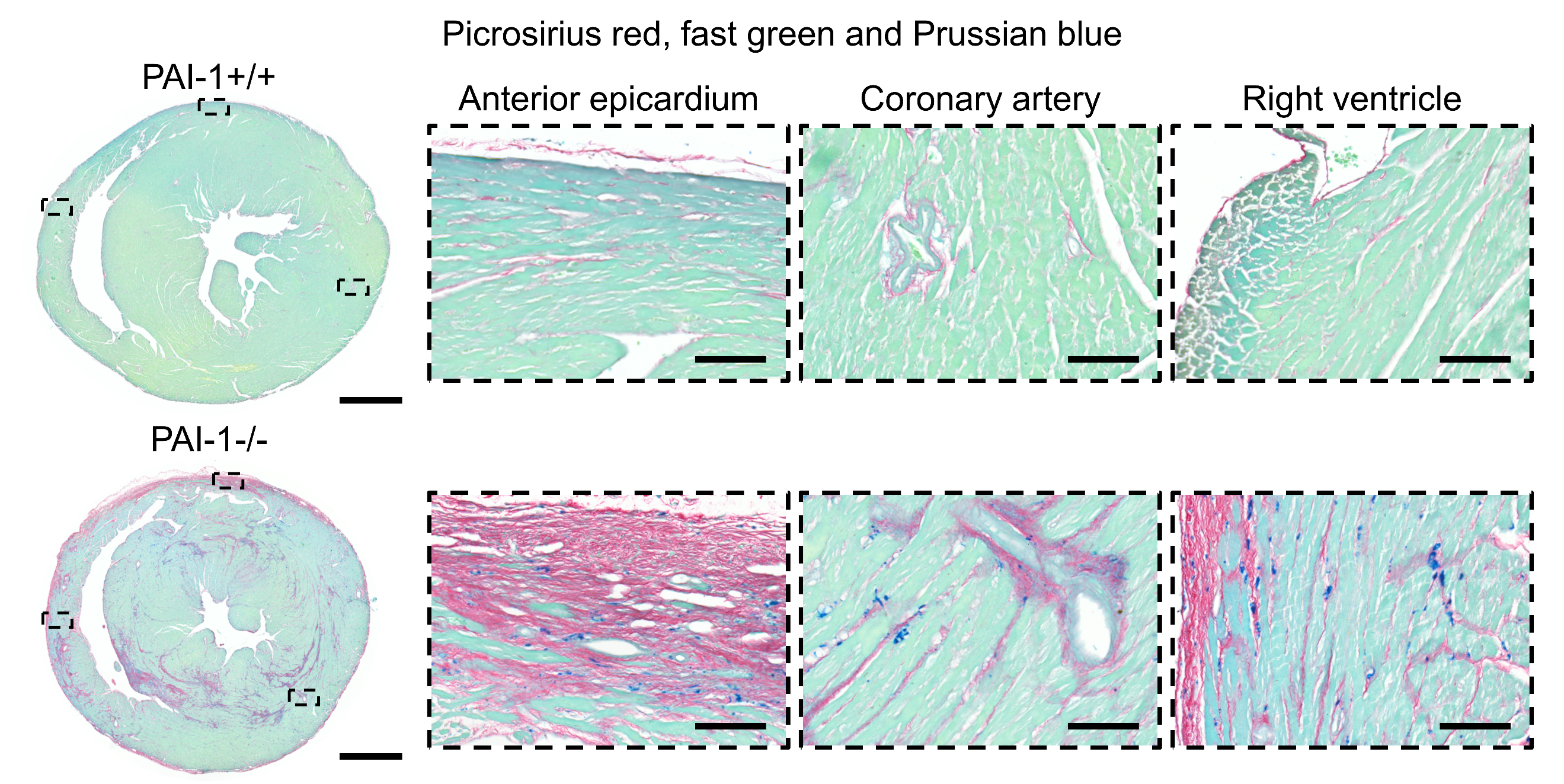
**Supplemental Figure 8.** **Ferric iron deposits were associated with fibrosis in PAI-1 deficient mice after 4 weeks of NE infusion.** Representative picrosirius red, fast green, and Prussian blue co-staining of mice infused with NE (5.6 mg/kg/day) for 4 weeks. Representative images are from male mice. Scale bars = 1,000 µm and 100 µm for whole section and high-magnification images, respectively. Abbreviations: NE: norepinephrine, PAI-1: plasminogen activator inhibitor-1.

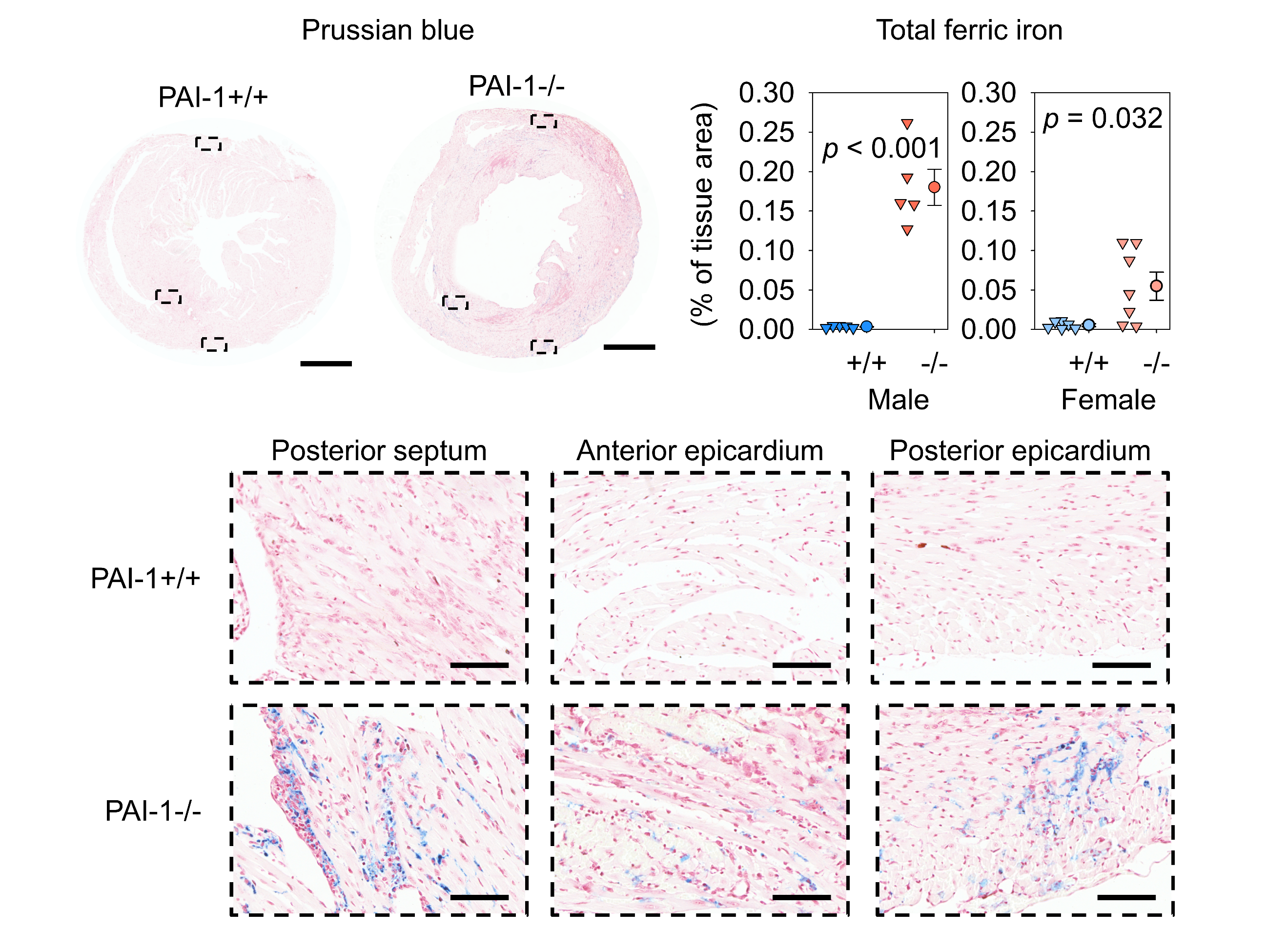
 **Supplemental Figure 9.** **PAI-1 deficiency induced ferric iron deposition after 1 week of AngII infusion.** Representative Prussian blue staining and ferric iron quantification of mice infused with AngII (1,000 ng/kg/min) for 1 week. Representative images are from male mice. Scale bars = 1,000 µm and 100 µm for whole section and high-magnification images, respectively. Abbreviations: AngII: angiotensin II, PAI-1: plasminogen activator inhibitor-1.

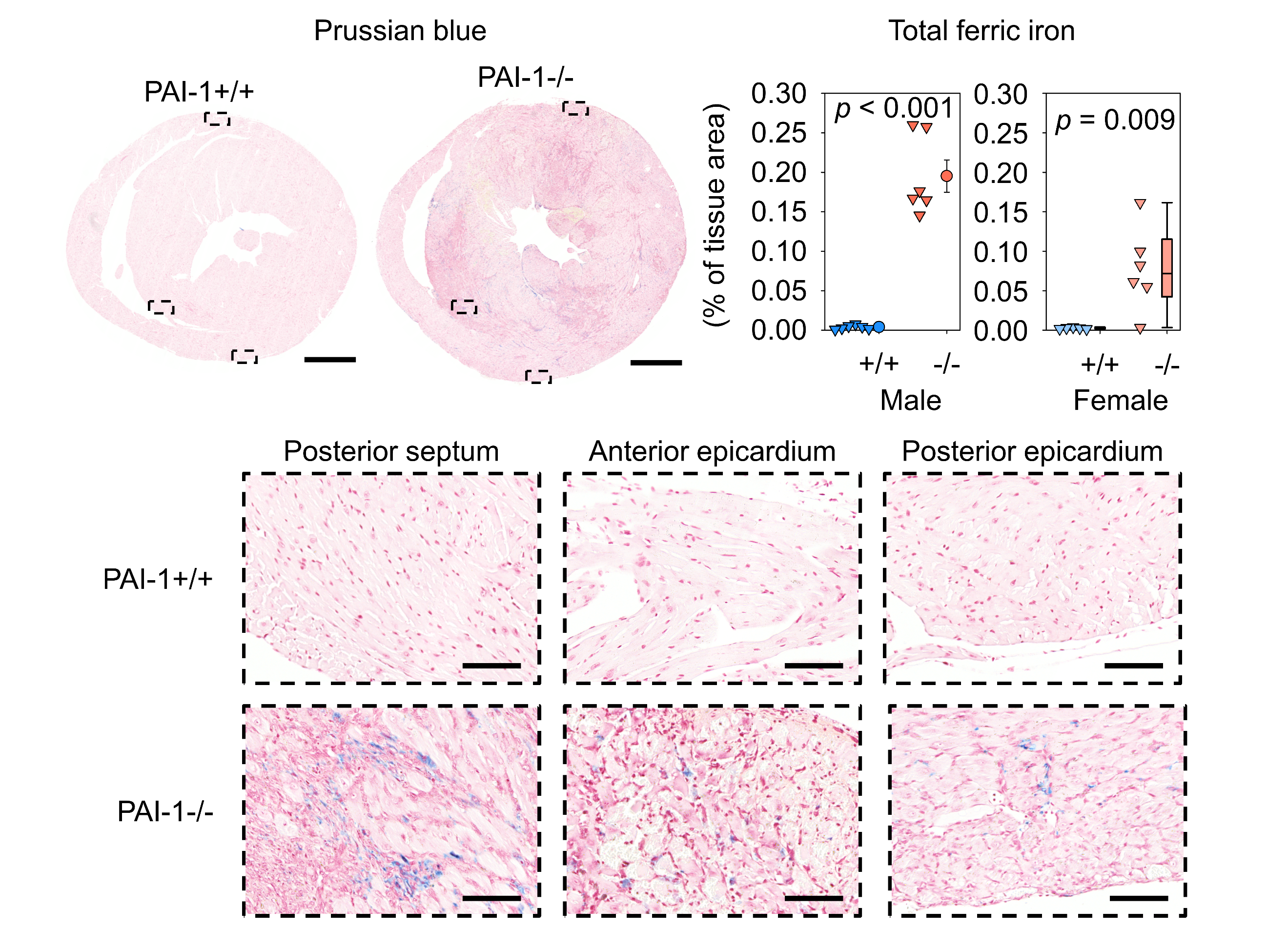
**Supplemental Figure 10.** **PAI-1 deficiency induced ferric iron deposition after 1 week of NE infusion.** Representative Prussian blue staining and ferric iron quantification of mice infused with NE (5.6 mg/kg/day) for 1 week. Representative images are from male mice. Scale bars = 1,000 µm and 100 µm for whole section and high-magnification images, respectively. Abbreviations: NE: norepinephrine, PAI-1: plasminogen activator inhibitor-1.

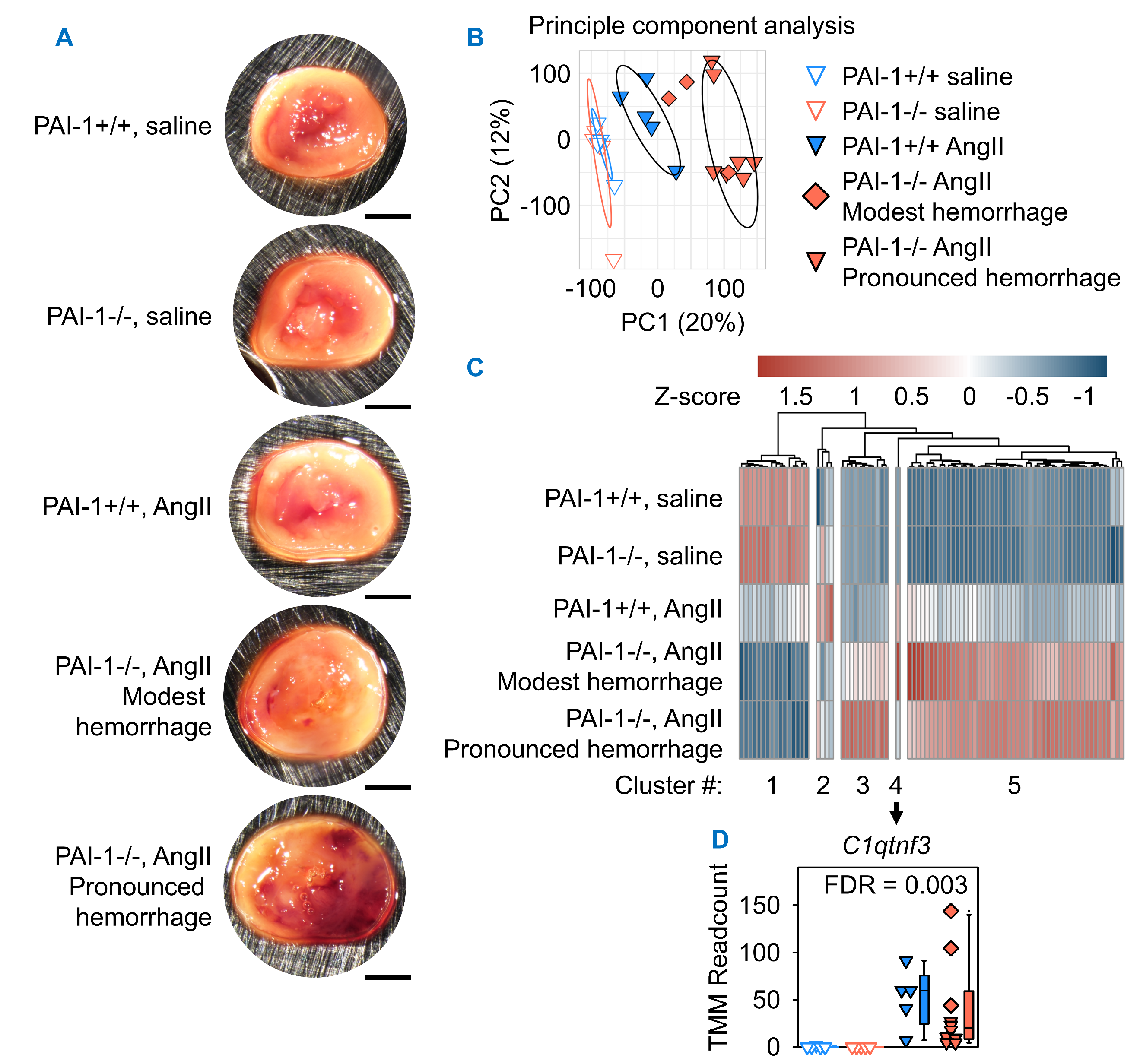

**Supplemental Figure 11. PAI-1 deficiency augmented AngII-induced cardiac transcriptomic alterations after 1 week of infusion.** Male PAI-1+/+ and -/- littermates were infused with saline or AngII (1,000 ng/kg/min) for 1 week. (**A**) Representative images of mid-ventricular heart sections used for bulk RNA-sequencing. Scale bar = 1,000 µm. (**B**) Principal component analysis of heart transcriptomes. Ellipses represent 75% confidence intervals. (**C**) Z-scored heatmap of all genes differentially expressed by interaction analysis of genotype and infusion controlling for the presence of pronounced hemorrhage. (**D**) Normalized read count expression of *C1qtnf3*, the sole transcript in cluster #4. Legend applies to (**B**) and (**D**). *n* = 5-10 mice/group. Abbreviations: PAI-1: plasminogen activator inhibitor-1, AngII: angiotensin II, PC: principal component, TMM: trimmed mean of M-values, FDR: false discovery rate.

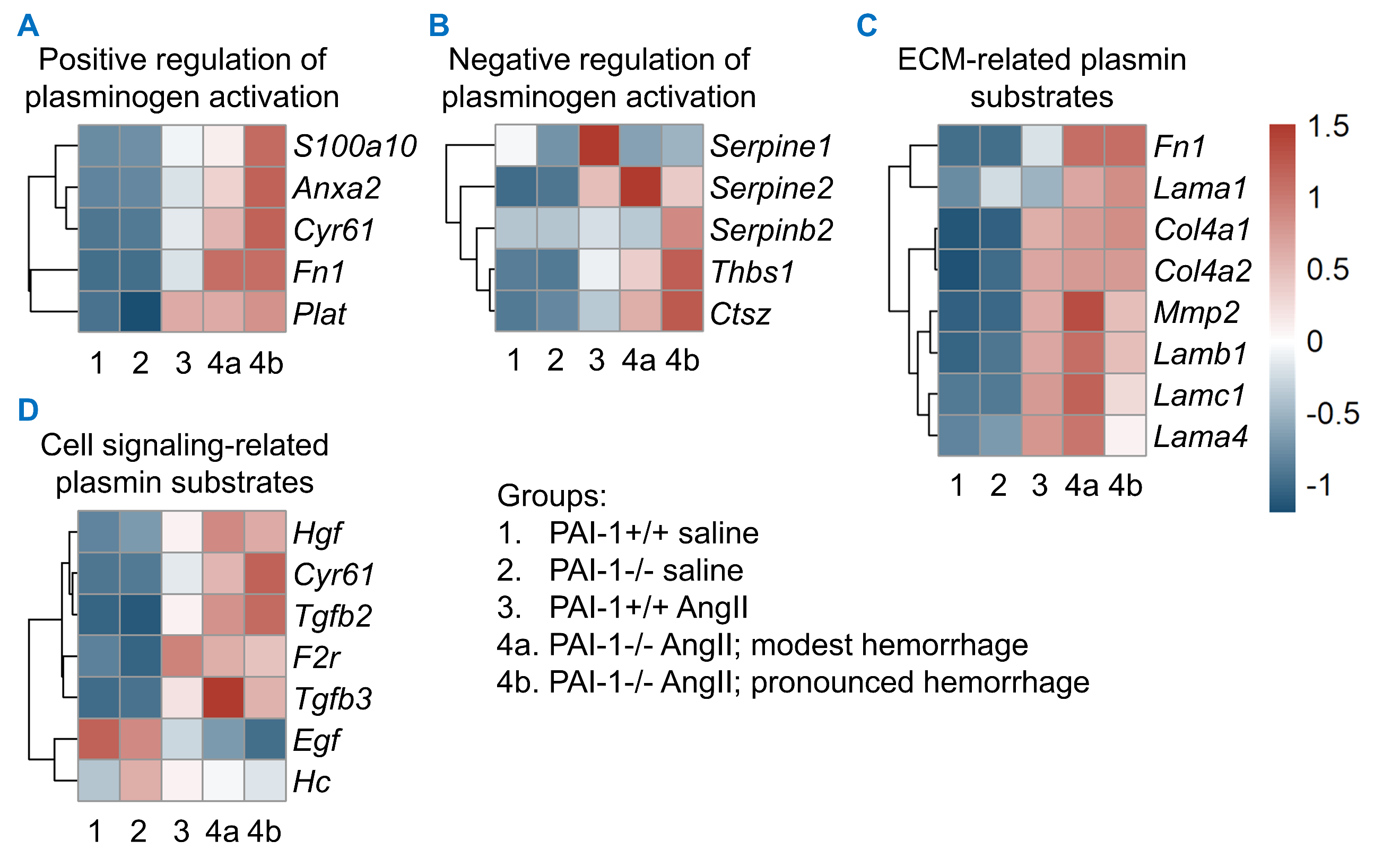
**Supplemental Figure 12. AngII infusion altered plasmin(ogen)-related cardiac transcripts after 1 week of infusion.** Male PAI-1+/+ and -/- littermates were infused with saline or AngII (1,000 ng/kg/min) for 1 week. Z-scored heatmaps of genes differentially expressed by AngII infusion analysis relating to positive (**A**) and negative regulation of plasminogen activation (**B**), and plasmin substrates related to extracellular matrix homeostasis (**C**) and cell-signaling (**D**). *n* = 5-10 mice/group. Abbreviations: PAI-1: plasminogen activator inhibitor-1, AngII: angiotensin II.

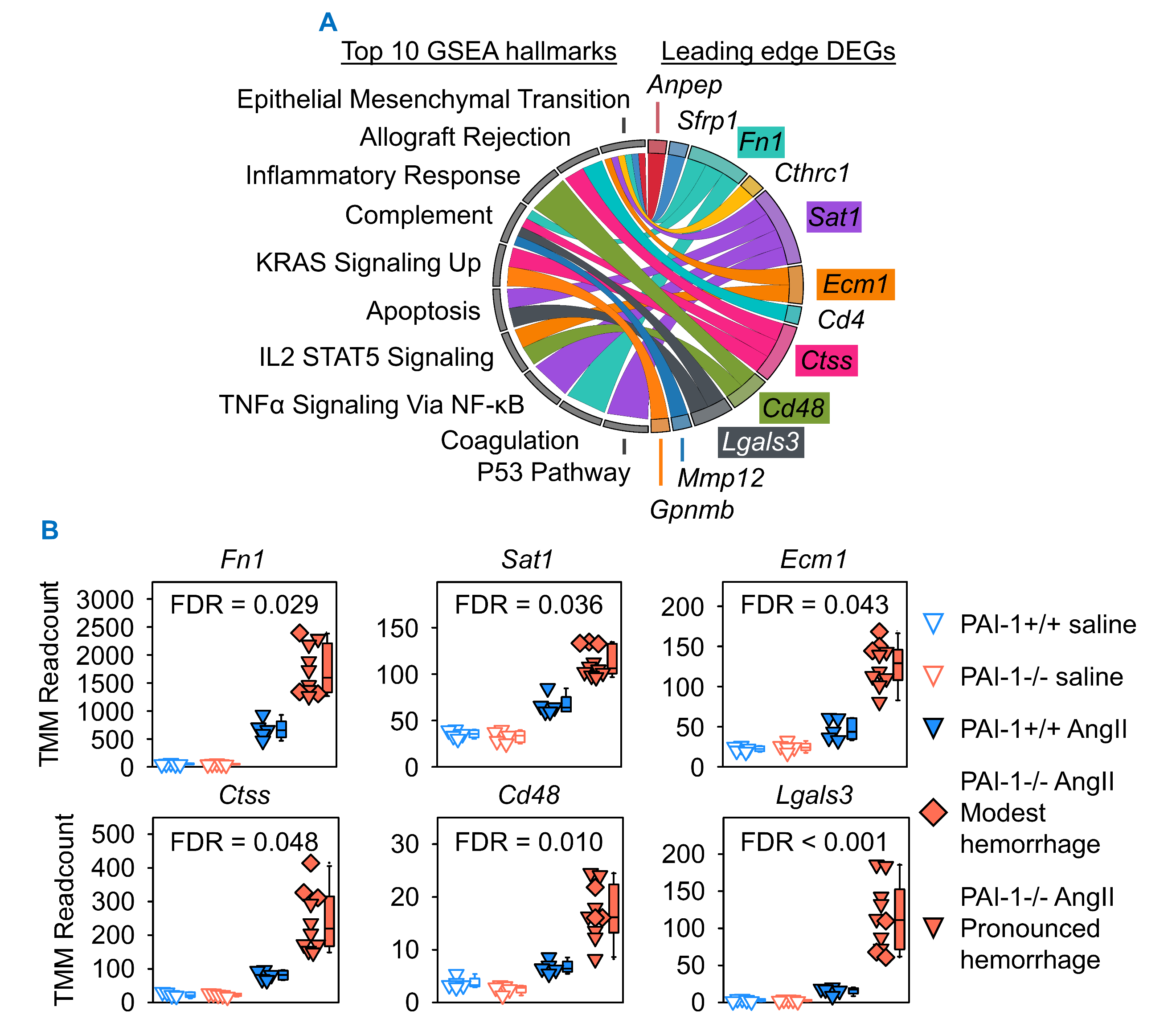

**Supplemental Figure 13. PAI-1 deficiency increased transcripts related to proteolysis and fibrosis independently of hemorrhage presence after 1 week of AngII infusion**. (**A**) Chord graph representing the top 10 GSEA hallmark gene sets and interaction DEGs present within the leading edge of corresponding Hallmark gene sets. (**B**) Normalized read count expression of interaction DEGs present in the leading edge of at least two top-10 GSEA Hallmark gene sets. *n* = 5-10 mice/group. Abbreviations: AngII: angiotensin II, PAI-1: plasminogen activator inhibitor-1, GSEA: gene set enrichment analysis, DEG: differentially expressed gene, TMM: trimmed mean of M-values, FDR: false discovery rate.

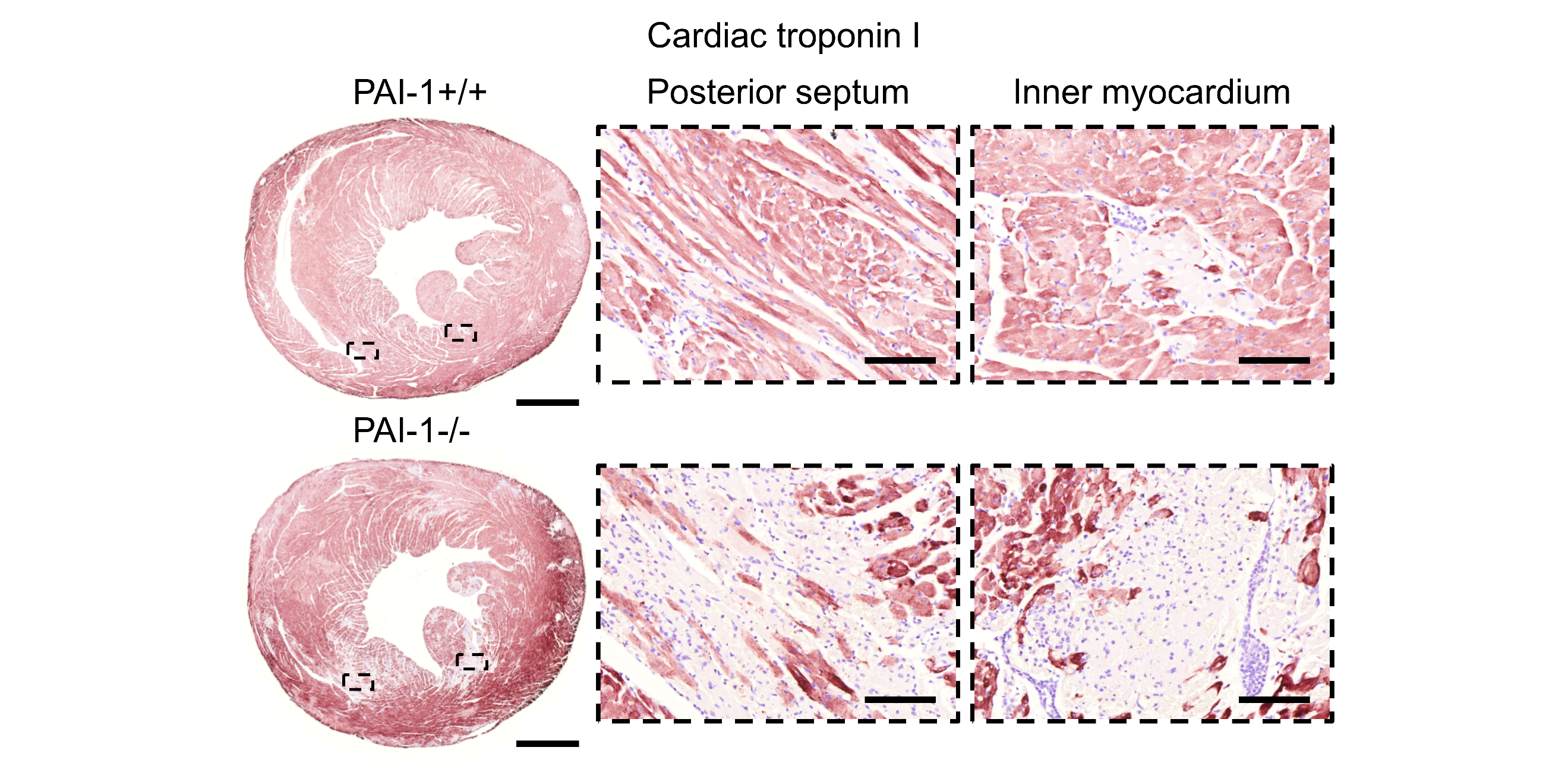

**Supplemental Figure 14.** **PAI-1 deficiency induced cardiomyocyte injury after 1 day of AngII infusion.** Male PAI-1+/+ and -/- littermates were infused with AngII (1,000 ng/kg/min) for 1 day. Representative cardiac troponin I immuno-histochemistry in mid-ventricular heart sections. Scale bars = 1,000 µm and 100 µm for whole section and high-magnification images, respectively. Abbreviations: AngII: angiotensin II, PAI-1: plasminogen activator inhibitor-1.

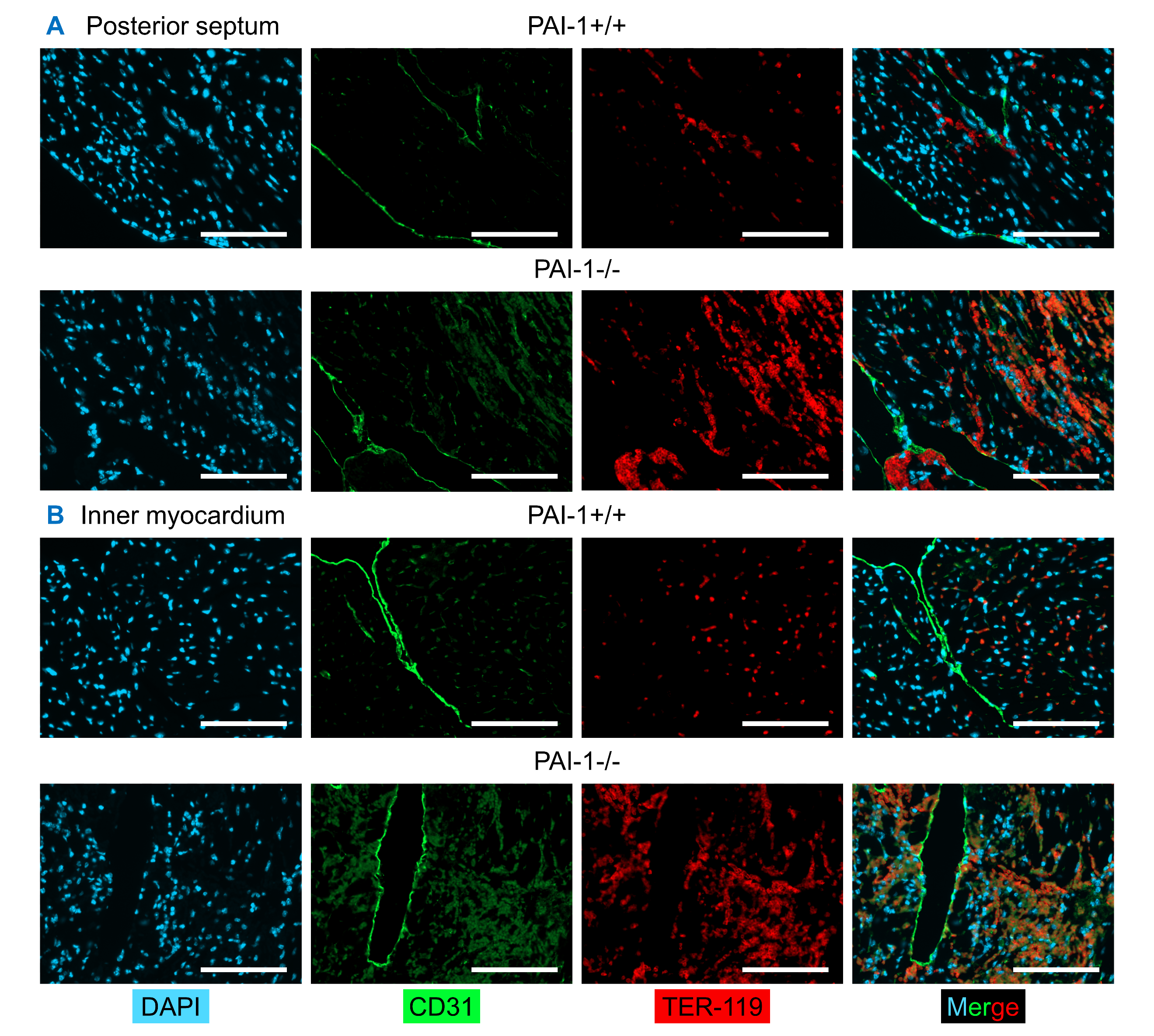

**Supplemental Figure 15.** **PAI-1 deficiency induced erythrocyte extravasation after 1 day of AngII infusion.** Male PAI-1+/+ and -/- littermates were infused with AngII (1,000 ng/kg/min) for 1 day. (**A-B**) Representative co-immunofluorescent staining for DAPI, CD31, and TER-119 in (**A**) the posterior septum and (**B**) the inner myocardium of mid-ventricular heart sections. Scale bars = 100 µm. Abbreviations: AngII: angiotensin II, PAI-1: plasminogen activator inhibitor-1, DAPI: 4',6-diamidino-2-phenylindole.

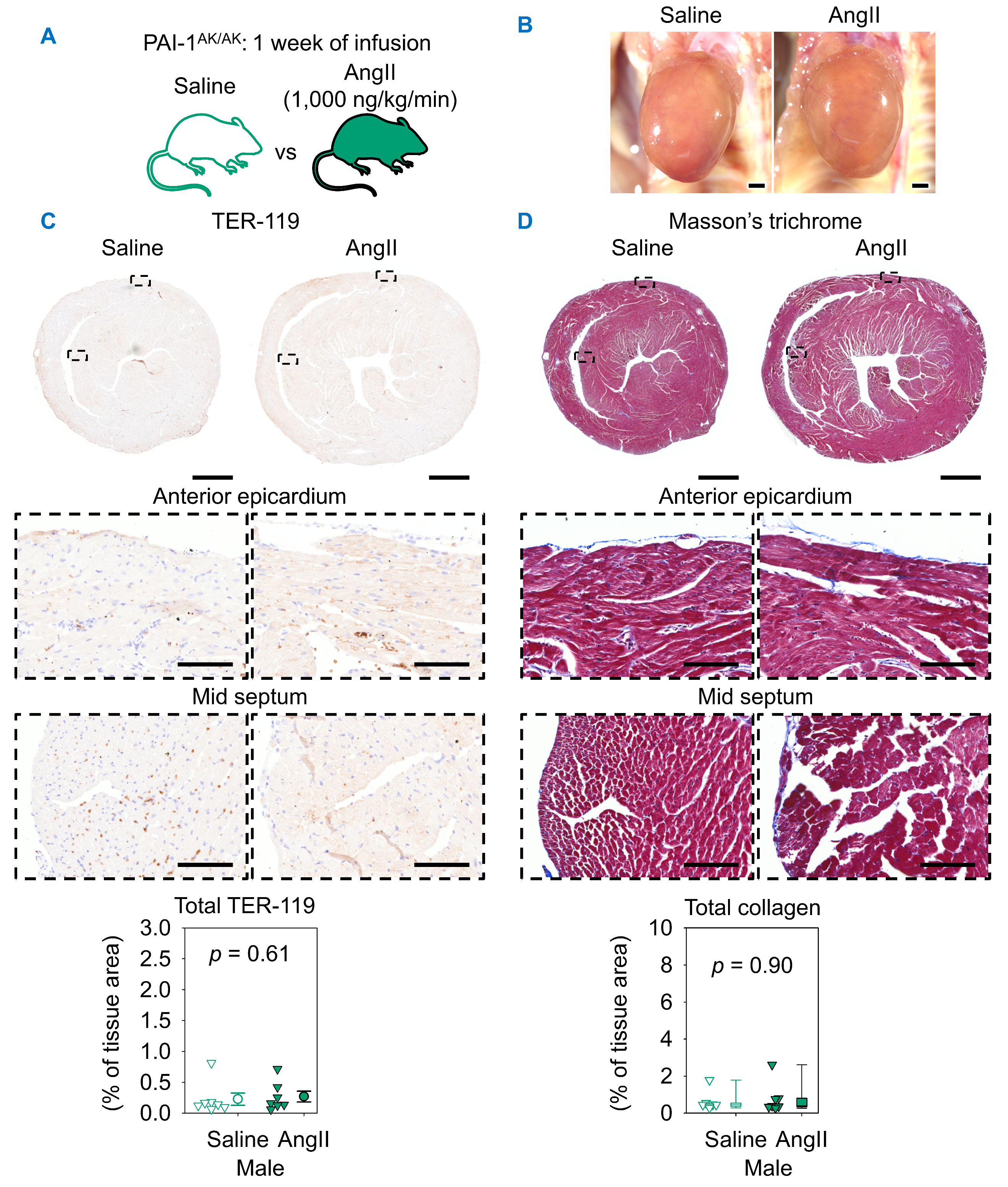

**Supplemental Figure 16. Mutational disruption of the somatomedin B-binding domain of PAI-1 did not contribute to AngII-induced cardiac injury.** (**A**) Male PAI-1AK/AK littermates were infused with saline or AngII (1,000 ng/kg/min) for 1 week. (**B**) Representative *in situ* heart images. (**C**) Representative TER-119 immunostaining and quantification of mid-ventricular hearts. Comparison was made by Student’s *t* test after log transformation. (**D**) Masson’s trichrome staining and collagen quantification of mid-ventricular hearts. Comparison was made by Mann-Whitney U test. *n* = 7 mice/group. Scale bars = 1,000 µm and 100 µm for whole section and high-magnification images, respectively. Abbreviations: AngII: angiotensin II, PAI-1: plasminogen activator inhibitor-1.

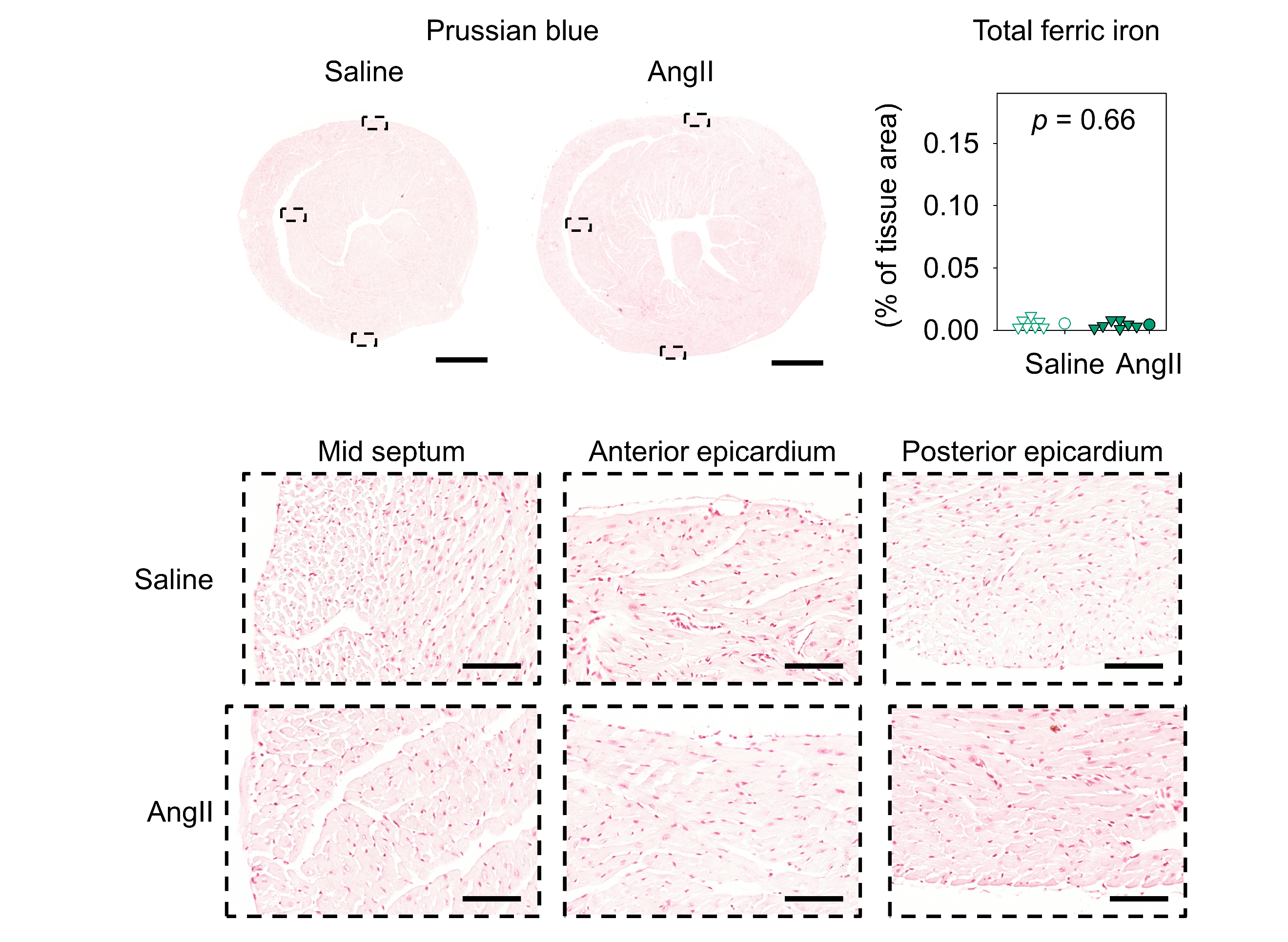
**Supplemental Figure 17. Mutational disruption of the somatomedin B-binding domain of PAI-1 did not contribute to AngII-induced ferric iron deposition.** Representative Prussian blue staining and ferric iron quantification of male PAI-1AK/AK mice infused with saline or AngII (1,000 ng/kg/min) for 1 week. Scale bars = 1,000 µm and 100 µm for whole section and high-magnification images, respectively. Abbreviations: AngII: angiotensin II, PAI-1: plasminogen activator inhibitor-1.

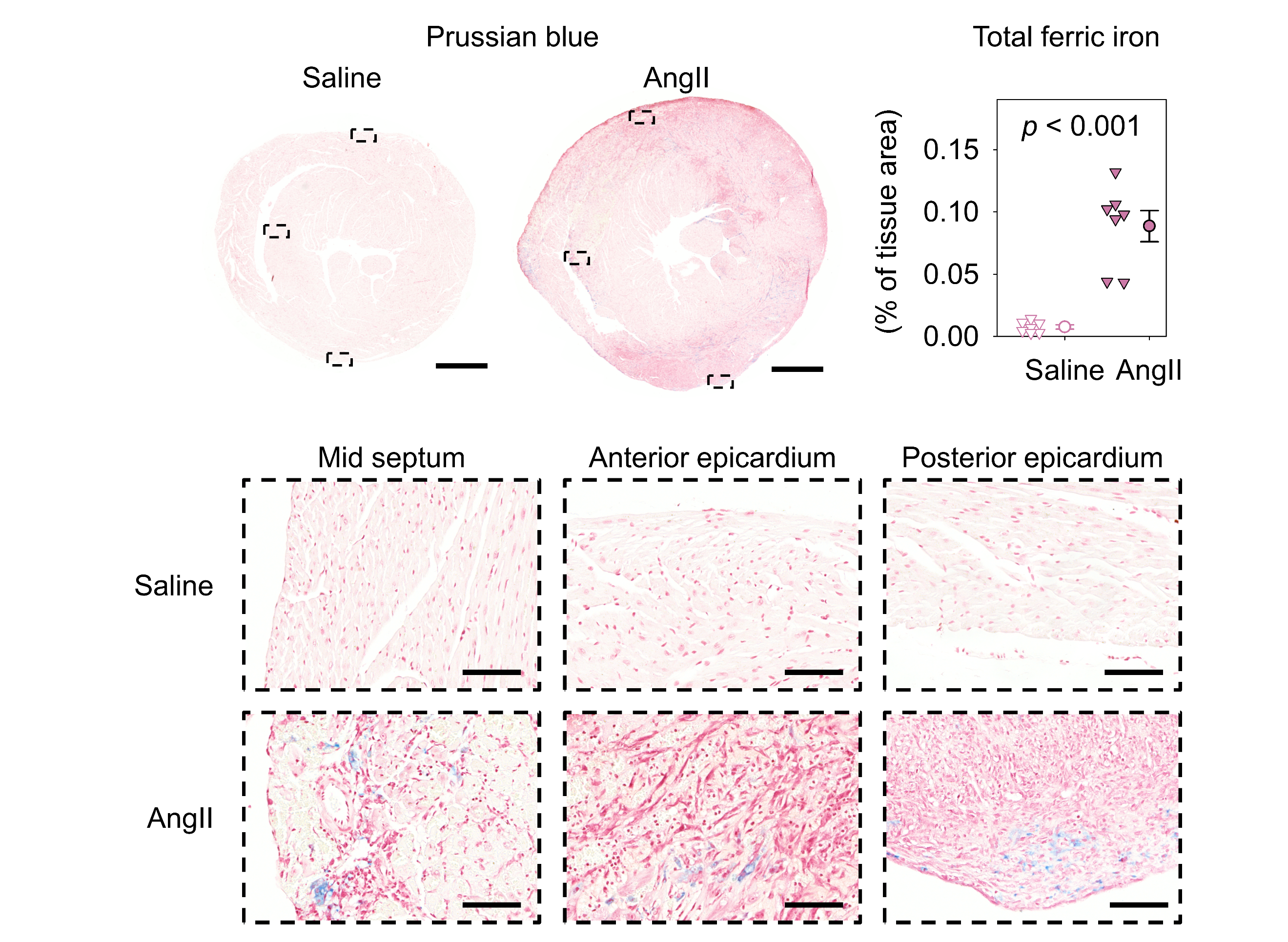
**Supplemental Figure 18. Mutational disruption of the reactive center loop domain of PAI-1 contributed to AngII-induced ferric iron deposition.** Representative Prussian blue staining and ferric iron quantification of male PAI-1Ala/Ala mice infused with saline or AngII (1,000 ng/kg/min) for 1 week. Scale bars = 1,000 µm and 100 µm for whole section and high-magnification images, respectively. Abbreviations: AngII: angiotensin II, PAI-1: plasminogen activator inhibitor-1.

**MAJOR RESOURCES TABLES**

**Primer sequences for genotyping of PAI-1+/+, PAI-1+/-, and PAI-1-/-**

| **Gene**  **(Forward / Reverse)** | **Vendor or Source** | **Primer Sequence (5’-3’)** |
| --- | --- | --- |
| PAI-1 (forward) | Integrated DNA Technologies | CTGGGCAGTAACCCAAGAGA |
| PAI-1 (wild type reverse) | Integrated DNA Technologies | GTCGGTCGTCTAGACCCTTG |
| PAI-1 (mutant reverse) | Integrated DNA Technologies | TGGATGTGGAATGTGTGCGAG |

**Primer sequences and restriction enzyme for genotyping of PAI-1^AK/AK^**

| **Gene**  **(Forward / Reverse)** | **Vendor or Source** | **Primer Sequence (5’-3’)** |
| --- | --- | --- |
| PAI-1 AK (forward) | Integrated DNA Technologies | ACCCAGAGGTGCATGGTGAG |
| PAI-1 AK (reverse) | Integrated DNA Technologies | GTGTGGCTCAGTGGGAGAGT |
| **Restriction Enzyme / Buffer** | **Vendor or Source** | **Catalog Number** |
| XbaI | New England Biolabs | R0145S |
| rCutSmart Buffer | New England Biolabs | B6004S |

**Primer sequences and restriction enzyme for genotyping of PAI-1^Ala/Ala^**

| **Gene**  **(Forward / Reverse)** | **Vendor or Source** | **Primer Sequence (5’-3’)** |
| --- | --- | --- |
| PAI-1 Ala (forward) | Integrated DNA Technologies | GATCTCTTGGGAATCACTCCA |
| PAI-1 Ala (reverse) | Integrated DNA Technologies | GTCCCTAGGGGCTCAAGAAA |
| **Restriction Enzyme / Buffer** | **Vendor or Source** | **Catalog Number** |
| NcoI | New England Biolabs | R3193S |
| rCutSmart Buffer | New England Biolabs | B6004S |

**Blocking reagents for immunostaining**

| **Reagent** | **Vendor** | **Catalog #** |
| --- | --- | --- |
| Normal goat serum | Vector laboratories | 30024 |
| Normal horse serum | Vector laboratories | 30022 |

**Primary antibodies for immunostaining**

| **Antibody** | **Vendor** | **Catalog #** | **Working concentration** |
| --- | --- | --- | --- |
| Rat anti-TER-119 | Thermo Scientific | 15-5921-82 | 0.3 µg/mL |
| Rabbit anti-cardiac troponin I | abcam | ab47003 | 1.0 µg/mL |
| Goat anti-CD31 | R&D Systems | AF3628 | 17.8 µg/mL |

**Secondary antibodies for immunostaining**

| **Antibody** | **Vendor** | **Catalog #** | **Working concentration** |
| --- | --- | --- | --- |
| ImmPRESS HRP goat anti-rat IgG | Vector Laboratories | 30033 | as supplied |
| ImmPRESS HRP goat anti-rabbit IgG | Vector Laboratories | 30125 | as supplied |
| Alexa-647 donkey anti-rat IgG | abcam | ab150155 | 2.0 µg/mL |
| Alexa-488 donkey anti-goat IgG | abcam | ab150129 | 2.0 µg/mL |

**RGB thresholds for histology and immunostaining quantification**

| **Stain** | **Feature** | **Thresholds** | | |
| --- | --- | --- | --- | --- |
|  |  | **Red** | **Green** | **Blue** |
| Masson’s trichrome | Collagen | 0–180 | 0–190 | 200–255 |
|  | Background | 220–255 | 200–255 | 215–255 |
| Prussian blue | Ferric iron | 30–150 | 40–160 | 70–240 |
|  | Background | 250–255 | 250–255 | 250–255 |
| TER-119 | Erythrocytes | 85–200 | 40–140 | 10–110 |
|  | Background | 250–255 | 250–255 | 250–255 |

**Figure 1**

| **Inclusion Criteria** | | | **Exclusion Criteria** | | | |
| --- | --- | --- | --- | --- | --- | --- |
| - Male and female - 8-14 weeks of age - PAI-1+/+ and PAI-1-/- | | | - Death prior to endpoint - Euthanasia prior to endpoint for humane reasons as based on IACUC regulations - Heterozygous genotype determined at endpoint verification - Plasma renin concentration (PRC) > 10 ng/ml/hour at endpoint | | | |
| **Groups** | **Sex** | **Age (weeks)** | | **Number (prior to experiment)** | **Number (termination)** | **Exclusions** |
| PAI-1+/+, AngII for 4 weeks | M | 8-14 | | 14 | 12 | (*n* = 2) death prior to endpoint |
| PAI-1-/-, AngII for 4 weeks | M | 8-14 | | 10 | 9 | (*n* = 1) PRC > 10 ng/ml/hour at endpoint |
| PAI-1+/+, AngII for 4 weeks | F | 8-14 | | 11 | 11 | none |
| PAI-1-/-, AngII for 4 weeks | F | 8-14 | | 22 | 16 | (*n* = 2) heterozygous genotype at endpoint; (*n* = 4) PRC > 10 ng/ml/hour at endpoint |

**Figure 2 and Supplemental Figure 1**

| **Inclusion/Exclusion Criteria** | | | | | |
| --- | --- | --- | --- | --- | --- |
| Inclusion and exclusion criteria are identical to those in Figure 1, with the following additional inclusion criteria:   - Random selection within groups when no gross cardiac pathology is present - Selection representative of the full range of observed gross cardiac pathology when present | | | | | |
| **Groups** | **Sex** | **Age (weeks)** | **Number (termination)** | **Number (histology)** | **Inclusion** |
| PAI-1+/+, AngII for 4 weeks | M | 8-14 | 12 | 8 | Random selection |
| PAI-1-/-, AngII for 4 weeks | M | 8-14 | 9 | 7 | Full range of gross pathology |
| PAI-1+/+, AngII for 4 weeks | F | 8-14 | 11 | 8 | Random selection |
| PAI-1-/-, AngII for 4 weeks | F | 8-14 | 16 | 9 | Full range of gross pathology |

Refer to Figure 1 table for the number of mice prior to experiment initiation and for exclusions.

**Figure 3 and Supplemental Figure 7**

| **Inclusion Criteria** | | | | | **Exclusion Criteria** | |
| --- | --- | --- | --- | --- | --- | --- |
| - Male and female - 10-12 weeks of age - PAI-1+/+ and PAI-1-/- - Random selection within groups when no gross cardiac pathology is present - Selection representative of the full range of observed gross cardiac pathology when present | | | | | - Death prior to endpoint - Euthanasia prior to endpoint for humane reasons as based on IACUC regulations - Heterozygous genotype determined at endpoint verification | |
| **Groups** | **Sex** | **Age (weeks)** | **Number (prior to experiment)** | **Number (histology)** | | **Inclusion / Exclusion** |
| PAI-1+/+, NE for 4 weeks | M | 11-12 | 7 | 5 | | Exclusion: (*n* = 1) heterozygous genotype at endpoint; (*n* = 1) death prior to endpoint  Inclusion: random selection |
| PAI-1-/-, NE for 4 weeks | M | 11-12 | 10 | 8 | | Exclusion: none  Inclusion: full range of gross pathology |
| PAI-1+/+, NE for 4 weeks | F | 11-12 | 9 | 6 | | Exclusion: none  Inclusion: random selection |
| PAI-1-/-, NE for 4 weeks | F | 11-12 | 10 | 5 | | Exclusion: none  Inclusion: full range of gross pathology |

**Figure 4 and Supplemental Figure 9**

| **Inclusion Criteria** | | | | | **Exclusion Criteria** | |
| --- | --- | --- | --- | --- | --- | --- |
| - Male and female - 10-12 weeks of age - PAI-1+/+ and PAI-1-/- - Random selection within groups when no gross cardiac pathology is present - Selection representative of the full range of observed gross cardiac pathology when present | | | | | - Death prior to endpoint - Euthanasia prior to endpoint for humane reasons as based on IACUC regulations - Heterozygous genotype determined at endpoint verification - Plasma renin concentration (PRC) > 10 ng/ml/hour at endpoint | |
| **Groups** | **Sex** | **Age (weeks)** | **Number (prior to experiment)** | **Number (histology)** | | **Inclusion / Exclusion** |
| PAI-1+/+, AngII for 1 week | M | 11-12 | 12 | 6 | | Exclusion: (*n* = 2) heterozygous genotype at endpoint  Inclusion: random selection |
| PAI-1-/-, AngII for 1 week | M | 11-12 | 15 | 5 | | Exclusion: (*n* = 3) heterozygous genotype at endpoint; (*n* = 3) death prior to endpoint; (*n* = 3) PRC > 10 ng/ml/hour at endpoint  Inclusion: full range of gross pathology |
| PAI-1+/+, AngII for 1 week | F | 11-12 | 18 | 7 | | Exclusion: (*n* = 2) heterozygous genotype at endpoint  Inclusion: random selection |
| PAI-1-/-, AngII for 1 week | F | 11-12 | 16 | 6 | | Exclusion: (*n* = 1) heterozygous genotype at endpoint; (*n* = 1) death prior to endpoint  Inclusion: full range of gross pathology |

**Figure 5 and Supplemental Figure 10**

| **Inclusion Criteria** | | | | | **Exclusion Criteria** | |
| --- | --- | --- | --- | --- | --- | --- |
| - Male and female - 10-12 weeks of age - PAI-1+/+ and PAI-1-/- - Random selection within groups when no gross cardiac pathology is present - Selection representative of the full range of observed gross cardiac pathology when present | | | | | - Death prior to endpoint - Euthanasia prior to endpoint for humane reasons as based on IACUC regulations - Heterozygous genotype determined at endpoint verification | |
| **Groups** | **Sex** | **Age (weeks)** | **Number (prior to experiment)** | **Number (histology)** | | **Inclusion / Exclusion** |
| PAI-1+/+, NE for 1 week | M | 11-12 | 10 | 6 | | Exclusion: (*n* = 1) heterozygous genotype at endpoint  Inclusion: random selection |
| PAI-1-/-, NE for 1 week | M | 11-12 | 9 | 6 | | Exclusion: none  Inclusion: full range of gross pathology |
| PAI-1+/+, NE for 1 week | F | 11-12 | 5 | 5 | | Exclusion: none  Inclusion: all |
| PAI-1-/-, NE for 1 week | F | 11-12 | 10 | 6 | | Exclusion: none  Inclusion: full range of gross pathology |

**Figure 6**

| **Inclusion Criteria** | | | **Exclusion Criteria** | | | |
| --- | --- | --- | --- | --- | --- | --- |
| - Male - 10-12 weeks of age - PAI-1+/+ and PAI-1-/- | | | - Death prior to endpoint - Euthanasia prior to endpoint for humane reasons as based on IACUC regulations - Heterozygous genotype determined at endpoint verification | | | |
| **Groups** | **Sex** | **Age (weeks)** | | **Number (prior to experiment)** | **Number (termination)** | **Exclusions** |
| PAI-1+/+, AngII for 1 day | M | 9-12 | | 6 | 4 | (*n* = 2) heterozygous genotype at endpoint |
| PAI-1-/-, AngII for 1 day | M | 9-12 | | 7 | 6 | (*n* = 1) heterozygous genotype at endpoint |

**Figure 7 and Supplemental Figure 18**

| **Inclusion Criteria** | | | **Exclusion Criteria** | | | |
| --- | --- | --- | --- | --- | --- | --- |
| - Male - 8-14 weeks of age - PAI-1Ala/Ala | | | - Death prior to endpoint - Euthanasia prior to endpoint for humane reasons as based on IACUC regulations - Endpoint verification genotype result different than weaning genotype | | | |
| **Groups** | **Sex** | **Age (weeks)** | | **Number (prior to experiment)** | **Number (termination)** | **Exclusions** |
| PAI-1Ala/Ala, saline for 1 week | M | 12 | | 7 | 7 | None |
| PAI-1Ala/Ala, AngII for 1 week | M | 12 | | 7 | 7 | None |

**Supplemental Figure 6**

| **Inclusion Criteria** | | | **Exclusion Criteria** | | | |
| --- | --- | --- | --- | --- | --- | --- |
| - Male and female - 10-12 weeks of age - PAI-1+/+ and PAI-1-/- | | | - Death prior to endpoint - Euthanasia prior to endpoint for humane reasons as based on IACUC regulations - Heterozygous genotype determined at endpoint verification | | | |
| **Groups** | **Sex** | **Age (weeks)** | | **Number (prior to experiment)** | **Number (termination)** | **Exclusions** |
| PAI-1+/+, saline for 4 weeks | M | 10-12 | | 8 | 7 | (*n* = 1) heterozygous genotype at endpoint |
| PAI-1-/-, saline for 4 weeks | M | 10-12 | | 9 | 8 | (*n* = 1) death prior to endpoint |
| PAI-1+/+, AngII for 4 weeks | M | 10-12 | | 10 | 8 | (*n* = 1) heterozygous genotype at endpoint; (*n* = 1) death prior to endpoint |
| PAI-1-/-, AngII for 4 weeks | M | 10-12 | | 11 | 7 | (*n* = 1) heterozygous genotype at endpoint; (*n* = 3) death prior to endpoint |
| PAI-1+/+, saline for 4 weeks | F | 10-12 | | 6 | 4 | (*n* = 2) heterozygous genotype at endpoint |
| PAI-1-/-, saline for 4 weeks | F | 10-12 | | 11 | 10 | (*n* = 1) heterozygous genotype at endpoint |
| PAI-1+/+, AngII for 4 weeks | F | 10-12 | | 7 | 6 | (*n* = 1) death prior to endpoint |
| PAI-1-/-, AngII for 4 weeks | F | 10-12 | | 11 | 9 | (*n* = 1) heterozygous genotype at endpoint; (*n* = 1) death prior to endpoint |

**Supplemental Figure 11-13**

| **Inclusion Criteria** | | | | **Exclusion Criteria** | |
| --- | --- | --- | --- | --- | --- |
| - Male and female - 10-12 weeks of age - PAI-1+/+ and PAI-1-/- - Within AngII-infused PAI-1-/-:   visible hemorrhage < 5% and ≥ 5% of tissue area = modest and pronounced hemorrhage, respectively | | | | - Death prior to endpoint - Euthanasia prior to endpoint for humane reasons as based on IACUC regulations - Heterozygous genotype determined at endpoint verification - RNA integrity number (RIN) < 6.8 | |
| **Groups** | **Sex** | **Age (weeks)** | **Number (prior to experiment)** | **Number (termination)** | **Exclusion** |
| PAI-1+/+, saline for 1 week | M | 8-12 | 5 | 5 | none |
| PAI-1-/-, saline for 1 week | M | 8-12 | 6 | 5 | (*n* = 1) RIN < 6.8 |
| PAI-1+/+, AngII for 1 week | M | 8-12 | 5 | 5 | none |
| PAI-1-/-, AngII for 1 week | M | 8-12 | 11 | 10 | (*n* = 1) death prior to endpoint |

**Supplemental Figures 16-17**

| **Inclusion Criteria** | | | **Exclusion Criteria** | | | |
| --- | --- | --- | --- | --- | --- | --- |
| - Male - 8-14 weeks of age - PAI-1AK/AK | | | - Death prior to endpoint - Euthanasia prior to endpoint for humane reasons as based on IACUC regulations - Endpoint verification genotype result different than weaning genotype | | | |
| **Groups** | **Sex** | **Age (weeks)** | | **Number (prior to experiment)** | **Number (termination)** | **Exclusions** |
| PAI-1AK/AK, saline for 1 week | M | 8-14 | | 7 | 7 | None |
| PAI-1AK/AK, AngII for 1 week | M | 8-14 | | 7 | 7 | None |

**ARRIVE Guidelines Checklists**

| **Item** | **Application** |
| --- | --- |
| Ethics | Approved by the University of Kentucky IACUC (2018-2967) |
| Sex | For 1 and 4-week infusion pathological studies in PAI-1+/+ and PAI-1-/- mice, both male and female animals were examined, and similar findings were reported for both sexes. Since no apparent sexual dimorphism was observed in AngII or NE-induced pathology, only male mice were examined for the 1-week bulk RNA sequencing and 1-day AngII infusion studies in PAI-1+/+ and PAI-1-/- mice, as well as the 1-week infusion studies in PAI-1Ala/Ala and PAI-1AK/AK mice. |
| Inclusion / Exclusion criteria | Criteria were determined *a priori* and are listed for individual experiments within the Major Resources Tables. |
| Sample size | Described in each figure legend. |
| Sample size calculation | Figure 1: *n* = 9/group were calculated to detect a 0.3 ± 0.2 mm (mean ± SD) difference in aortic diameter (*t* test, α error = 0.05 and power = 0.8).  Figure 3: *n* = 7/group were calculated to detect a 6 ± 3.6% (mean ± SD) difference in cardiac collagen staining (*t* test, α error = 0.05 and power = 0.8).  Supplemental Figure 6: *n* = 9/group were calculated to detect a 20 ± 10% (mean ± SD) difference in ejection fraction (ANOVA, α error = 0.05 and power = 0.8).  Figure 7 and Supplemental Figures 16-18: *n* = 7/group were calculated to detect a 50 ± 30% (mean ± SD) difference in cardiac TER-119 staining (*t* test, α error = 0.05 and power = 0.8).  Sample size calculations were not performed for the remaining figures. |
| Endpoint | Figure 1: ascending aortic diameters.  Figures 2-3: cardiac collagen and ferric iron.  Figures 4-5: cardiac collagen and TER-119.  Figure 6: cardiac TER-119.  Figure 7: cardiac collagen and TER-119.  Supplemental Figure 6: ejection fraction.  Supplemental Figure 16: cardiac collagen and TER-119. |
| Randomization | Mice were randomly assigned to cages at weaning by sex. After stratifying by age, mice were randomly assigned to study groups. Mice were terminated without predetermined order. |
| Blinding | Genotyping, measurement of ultrasound images and *in situ* aortas, and quantification of histology were performed by an investigator blinded to the study group information. RNA sequencing library generation was performed by a 3^rd^ party blinded to the study group information. All experimental data were verified by an independent investigator blinded to the study group information. |
| Statistical analysis | SigmaPlot (v16.0, SYSTAT Software Inc.) or R (v4.5.1). |
| Statistical method | Described in each figure legend. |
| Data availability | RNA sequencing data (raw FASTQ and aligned data) will be made publicly available at the Gene Expression Omnibus (GEO) repository. Numerical data that support the findings of this study are provided in the Supporting Data Values file. The data that support the findings of this study are also available from the corresponding authors on reasonable request. |
